# Activation of a Src-JNK pathway in unscheduled endocycling cells of the *Drosophila* wing disc induces a chronic wounding response

**DOI:** 10.1101/2025.03.12.642788

**Authors:** Yi-Ting Huang, Brian R. Calvi

## Abstract

The endocycle is a specialized cell cycle during which cells undergo repeated G / S phases to replicate DNA without division, leading to large polyploid cells. The transition from a mitotic cycle to an endocycle can be triggered by various stresses, which results in unscheduled, or induced endocycling cells (iECs). While iECs can be beneficial for wound healing, they can also be detrimental by impairing tissue growth or promoting cancer. However, the regulation of endocycling and its role in tissue growth remain poorly understood. Using the *Drosophila* wing disc as a model, we previously demonstrated that iEC growth is arrested through a Jun N-Terminal Kinase (JNK)-dependent, reversible senescence-like response. However, it remains unclear how JNK is activated in iECs and how iECs impact overall tissue structure. In this study, we performed a genetic screen and identified the Src42A-Shark-Slpr pathway as an upstream regulator of JNK in iECs, leading to their senescence-like arrest. We found that tissues recognize iECs as wounds, releasing wound-related signals that induce a JNK-dependent developmental delay. Similar to wound closure, this response triggers Src-JNK-mediated actomyosin remodeling, yet iECs persist rather than being eliminated. Our findings suggest that the tissue response to iECs shares key signaling and cytoskeletal regulatory mechanisms with wound healing and dorsal closure, a developmental process during *Drosophila* embryogenesis. However, because iECs are retained within the tissue, they create a unique system that may serve as a model for studying chronic wounds and tumor progression.

**Article summary:** The effects of unscheduled endocycles on tissue growth remain unclear. To investigate this, we used *Drosophila* to induce a switch from the mitotic cycle to the endocycle and analyzed tissue responses at both the signaling and tissue structure levels. Surprisingly, tissues recognized endocycling cells as wounds, activating regeneration signals and remodeling tissue structure. However, because these cells resist apoptosis, they persist within the tissue without being cleared. This persistence disrupts normal healing, revealing the similarities between unscheduled endocycling cells and chronic wounds. Our system has the potential to serve as a novel model for studying chronic wound responses or tumorigenesis.

## Introduction

The regulation of tissue growth and homeostasis is essential for normal development and tissue function. Tissues grow through an increase in cell number, but can also grow through an increase in cell size. One way that cells grow in size is through the endocycle, which is a cell cycle variant that is composed of G / S phases without mitotic cell division. Over multiple G / S endocycles, cells increase in size (hypertrophy) and repeatedly duplicate their genome, resulting in large, polyploid cells (Ovrebo and Edgar 2018). Endocycles are a normal developmental growth program of specific tissues across the tree of life, including plants, insects, and mammals (Morris *et al*. 2024). In addition to developmentally programmed endocycles, cells can also initiate endocycles in response to various conditional signals (Herriage *et al*. 2024). In this study, we used the *Drosophila* wing disc as a model to define the impact of these induced endocycling cells (iECs) on tissue growth.

Cells can switch to unscheduled endocycles in response to various stresses, during aging, and after wounding (Ivanov *et al*. 2003; Losick *et al*. 2016; Ovrebo and Edgar 2018; Matsumoto *et al*. 2021; Huang *et al*. 2024b). In some contexts, iECs can be beneficial by facilitating wound healing and regeneration (Losick *et al*. 2013; Tamori and Deng 2013; Cao *et al*. 2017; Cohen *et al*. 2018; Ovrebo and Edgar 2018; Bailey *et al*. 2021; Kirillova *et al*. 2021; Chakraborty *et al*. 2023; Zuppo *et al*. 2023). In other contexts, however, they can inhibit tissue growth and function (Gonzalez-rosa *et al*. 2018; De Chiara *et al*. 2022; Herriage and Calvi 2024; Huang *et al*. 2024b). In addition, there is evidence that unscheduled endocycles can be detrimental to organismal health by contributing to cancer. These endocycling cancer cells are commonly called Polyploid Giant Cancer Cells (PGCCS) and have been reported in tumors of *Drosophila* and a wide variety of human cancers (Tamori and Deng 2013; Zack *et al*. 2013; Almeida Machado Costa *et al*. 2022; Liu *et al*. 2022; Herriage *et al*. 2024). Similar to iECs in *Drosophila*, human PGCCS are resistant to cell death, thereby contributing to cancer therapy resistance (Illidge *et al*. 2000; Hassel *et al*. 2014; Zhang *et al*. 2014; Mittal *et al*. 2017). Also similar between fly and human, PGCCs can resume error-prone polyploid divisions, which generates genetically-diverse pools of aneuploid daughter cells that have the potential to contribute to cancer progression (Puig *et al*. 2008; Erenpreisa *et al*. 2011; Hassel *et al*. 2014; Chen *et al*. 2016). Therefore, unscheduled endocycling cells can be either beneficial or harmful, but what determines these disparate effects is largely undefined.

We have been investigating the regulation of iECs and their effects on cell and tissue growth in *Drosophila* (Maqbool *et al*. 2010; Hassel *et al*. 2014; Qi and Calvi 2016; Rotelli *et al*. 2019a; Rotelli *et al*. 2019b; Herriage and Calvi 2024; Huang *et al*. 2024b). Through the genetic inhibition of mitosis, we can experimentally induce cells in any tissue to switch from mitotic cycles to unscheduled endocycles (Verghese and Su 2017; Herriage and Calvi 2024; Huang *et al*. 2024b). We previously reported that populations of iECs in the larval wing disc grow to different cell sizes and DNA ploidies before entering a senescent-like arrest (Huang *et al*. 2024b). Unlike other growth-challenged cells, these arrested iECs were resistant to apoptosis and not eliminated from the disc epithelium. Although senescent iECs stimulated compensatory proliferation of neighboring diploid cells, this was unable to regenerate a normal disc. We found that the iEC senescent arrest was induced by activation of c-Jun N-terminal kinase (JNK). JNK, encoded by the *basket* (*bsk*) gene in *Drosophila*, is part of a conserved kinase pathway that is activated by a variety of upstream stimuli to mediate a number of different downstream effects (Johnson and Nakamura 2007; La Marca and Richardson 2020; Tafesh-Edwards and Eleftherianos 2020). JNK is known to mediate dynamic tissue reorganization during developmental morphogenesis, wound healing, and regeneration, and, depending on context, can either inhibit or promote growth and oncogenesis (Schaeffer and Weber 1999; Enomoto *et al*. 2015; Smith-Bolton 2016; La Marca and Richardson 2020).

Although it is clear that activation of JNK restrains iEC growth, which upstream signals activate JNK in iECs and how this signaling affects tissue morphology is not known.

In this study, we found that a Src signaling pathway is required to activate JNK in wing disc iECs. Activation of this Src-JNK pathway induced a senescent-like arrest, gene expression, and a dynamic tissue reorganization that were all similar to wound healing response. Overall, our findings suggest that persistent unscheduled endocycling cells can be detrimental to tissue growth by inducing a chronic wounding response, with broader relevance to how PGCCs contribute to tumorigenesis.

## Materials & Methods

### *Drosophila* strains and genetics

Information about fly strains, genetics, and other information was obtained from FlyBase (Ozturk-Colak *et al*. 2024). Most fly strains were obtained from Bloomington *Drosophila* Stock Center, and were raised in standard Bloomington *Drosophila* stock center media (https://bdsc.indiana.edu/information/recipes/bloomfood.html) at 25°C. For GAL80^ts^ experiments, larvae were raised at 18°C then shifted to 29°C to activate GAL4 drivers. Fly strains obtained from Bloomington *Drosophila* Stock Center (BDSC, Bloomington, IN, USA) for the genetic screen are listed in Table S1. Strains that were used in individual panels are listed in Table S2.

### Immunofluorescence microscopy and quantification

Late 3^rd^ instar larvae were dissected in PBS (phosphate buffered saline), fixed in 6% formaldehyde (Avantor, Cat# MAL-5016-02), permeabilized with PBT (phosphate buffered saline with 0.1% Triton X-100), and blocked in 5 % Normal Goat Serum (Gibco, Cat#16210072) as previously described (Huang *et al*. 2024b). The following antibodies were used: anti-RFP (Takara, Cat#632496, RRID:AB_10013483) at 1:1000, anti-GFP (Invitrogen, Cat#A11120, RRID:AB_221568) at 1:1000, anti-GFP (Invitrogen, Cat#A11122, RRID:AB_221569) at 1:1000, anti-β-galactosidase (Abcam, Cat#ab9361, RRID:AB_307210) at 1:2000. For the F-actin labeling, tissues were incubated in fluorescent labeled phalloidin (Invitrogen, Cat#A22284) at 1:200 for 2 hours. The tissues were stained with DAPI (0.5 μg/ml) and imaged on a Leica SP8 confocal. All y-z images were processed in ImageJ software (RRID:SCR_003070). Fig. 2g was quantified by using Imaris software (RRID:SCR_007370). Cells were recognized by Imaris machine learning, and the ploidy fold change was quantified by the sum of DAPI intensity in the GAL4-expressing cells and was normalized to the GAL4-negative cells in the same disc. All videos were made and recorded by using Imaris software (RRID:SCR_007370).

### SA-β-GAL activity labeling

The larvae were dissected during late 3^rd^ instar in PBS, fixed in 4 % formaldehyde (Electron Microscopy Sciences, Cat#157-4-1L), washed with PBS plus 2.5 % Bovine Serum Albumin (BSA) (Fisher BioReagents, Cat#BP1605-100). β-GAL activity was detected by incubating tissues at 37 °C for 3 hours according to the manufacturer’s instructions (Invitrogen, Cat#C10850). The tissues were then washed with PBT for 3 times 10-min and stained with DAPI (0.5 μg/ml).

### Dihydroethidium (DHE) staining

The larvae were dissected at late 3^rd^ in PBS, and incubated with 30 uM dihydroethidium (DHE) (Invitrogen Molecular Probes, Cat#D11347) for 5 min in the dark, then washed with PBS for 3 times 10-min. The tissues were mounted with 50 % glycerol and imaged on a Leica SP8 confocal immediately.

### Statistical Analysis

Statistical analysis was performed using GraphPad Prism. The data in graph in Fig. 2g is shown as mean ± S.D. from at least three biological replicates, and a one-way ANOVA was used to calculate p values. All data shown is based upon at least three biological replicates.

## Results

### A candidate genetic screen to identify upstream activators of JNK in iECs

To investigate JNK activation in iECs and its effect on cell and tissue growth, we used the *Drosophila* wing disc as a model (Figure 1a). The wing disc is a sac-like structure composed of two layers of epithelial cells separated by a lumen; the disc proper epithelium (DP) and the peripodial epithelium (PE) (Auerbach 1936; Milner *et al*. 1984; Tripathi and Irvine 2022). The DP is fated to become adult structures and during larval development undergoes approximately 10 mitotic cycles to increase from ∼30 to ∼30,000 cells. Wing discs can regenerate after surgical or genetic ablation, and analysis of this regeneration has led to discovery of conserved mechanisms that regulate tissue growth and tumorigenesis from flies to humans (Vidal and Cagan 2006; Aegerter-Wilmsen *et al*. 2007; Neto-Silva *et al*. 2009; Hariharan 2015; Tamori *et al*. 2016; Morata and Calleja 2020; Morata 2021). Wing discs are also a premiere model for mechanisms of developmental patterning (Tripathi and Irvine 2022). Cells in the central wing pouch are fated to become adult wing blade, a ring of cells surrounding the pouch develop into the more proximal adult wing hinge, while the disc notum forms part of the adult thorax (Figure 1a) (Tripathi and Irvine 2022). Characteristic folds in the DP epithelium serve as morphological landmarks for these different regions.

**Figure 1.**
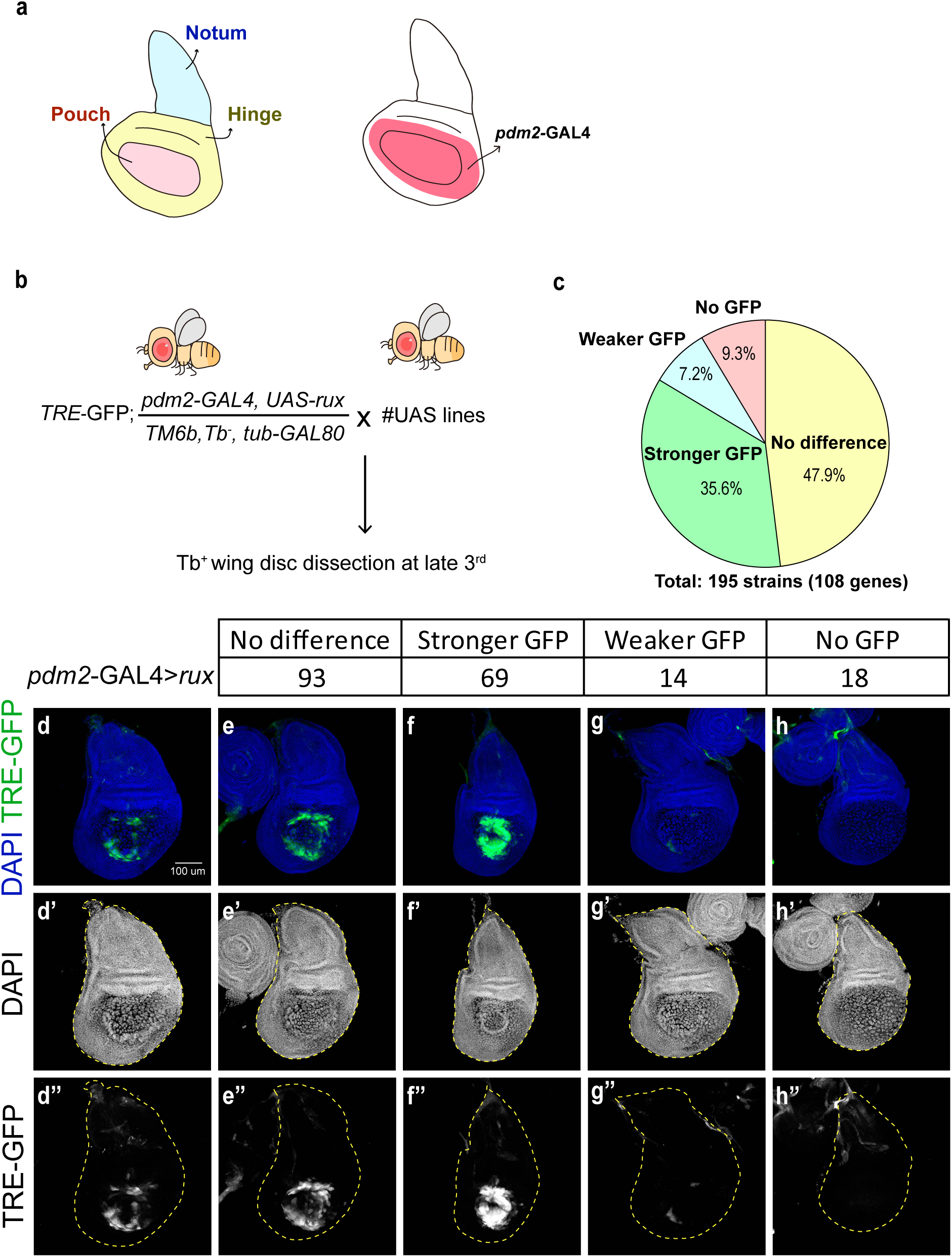
Genetic screen to identify upstream activators of JNK in iECs. (a) Schematic representation of a *Drosophila* wing disc. The wing disc is subdivided into the wing notum (blue), wing hinge (yellow), and wing pouch (pink). *Pdm2-*GAL4>GAL4 is expressed in the wing pouch and the inner wing hinge, highlighted in red. (b) Crossing scheme for the genetic screen. A fly strain with a JNK reporter, TRE-GFP, and *pdm2-*GAL4>*rux* was maintained over a balancer with the GAL4 repressor, GAL80. This strain was crossed to different UAS RNAi or other UAS strains and wing discs were dissected from the Tb+ larvae with GAL4 activity during late 3^rd^ instar to analyze TRE-GFP expression in iECs. (c-h’’) Summary of the results of genetic screen. A total of 195 strains representing 108 genes were analyzed and divided into four categories based on the GFP signal. The distribution of each category is summarized in (c) and the images of the representative strains of each category are shown in (d-h’’). Scale bar: 100 µm. Full strain list is available in Table S1.

We switched central pouch and surrounding proximal wing hinge into endocycles using *pdm2-GAL4* (Butler *et al*. 2003; Jory *et al*. 2012; Loker and Mann 2022) to express a GAL4-inducible transgene containing *roughex* (*rux*), a Cyclin A inhibitor (Figure 1a) (Jones and Moses 2004; Jenett *et al*. 2012). We had previously shown that these iECs grow in size and DNA content but then activate JNK and arrest (Huang *et al*. 2024b). We detected JNK activity using a reporter that has an artificial tetradecanoylphorbol acetate response element fused to EGFP (TRE-GFP) (Chatterjee and Bohmann 2012; Huang *et al*. 2024b). This reporter contains binding sites for the conserved AP-1 transcription factor, which is phosphorylated and activated by JNK (Angel and Karin 1991). A large diversity of stimuli and signaling pathways are known to activate JNK, and it remained unclear which of these upstream pathways activate JNK in iECs (La Marca and Richardson 2020). To address this question, we performed a candidate genetic screen. We selected strains with GAL4-inducible RNAi, dominant-negative, and constitutively active transgenes that alter the activity of genes that have been previously implicated in regulating the JNK pathway in other contexts (Table S1). We screened for the effect of these transgenes on TRE-GFP expression in wing iECs (Figure 1b). We screened a total of 195 strains representing 108 genes and divided results into four categories based on TRE-GFP expression: no change (93 strains, 47.9%), stronger GFP (69 strains, 35.6%), weaker GFP (14 strains, 7.2%), or no GFP (18 strains, 9.3%) (Figure 1c-h’’). A full strain and gene list for the screen is summarized in Table S1.

### The TNF pathway and ROS do not play major roles in JNK activation in iEC

Previous studies have shown that the Tumor necrosis factor (TNF) pathway is one of the major upstream activators of the JNK pathway (Figure S1a) (Igaki *et al*. 2002; Marchal *et al*. 2012; La Marca and Richardson 2020; Herrera and Bach 2021). However, knockdown of the single TNF ligand in *Drosophila*, *eiger (egr)* (Igaki *et al*. 2002; Zhang *et al*. 2024), or its receptor, *wengen (wgn)* (Kanda *et al*. 2002) did not alter TRE-GFP expression in iEC (Figure S1b-d’). Moreover, knockdown of downstream TNF pathway members *misshapen (msn)* (Treisman *et al*. 1997; Su *et al*. 1998), *TGF-β activated kinase 1 (tak1)* (Yamaguchi *et al*. 1995; Takatsu *et al*. 2000), *MAP kinase kinase 4 (Mkk4)* (Derijard *et al*. 1995; Lin *et al*. 1995; Han *et al*. 1998), or overexpression of a dominant negative form of the TNF pathway kinase *wallenda (wnd^K118^)* (Collins *et al*. 2006) also did not affect JNK activity in iEC (Figure S1e-h’). All of these strains have been shown to compromise TNF signaling and JNK activation in other contexts, yet none of them had a measurable effect on JNK activity in iECs, strongly suggesting that the TNF pathway does not play a major role in activating JNK during unscheduled endocycles.

Radical oxygen species (ROS) and hydrogen peroxide (H_2_O_2_) are also known to activate JNK pathway through Apoptotic signal-regulating kinase 1 (Ask1, a JNKKK). We assayed ROS and H_2_O_2_ using dihydroethidium (DHE) staining (Owusu-Ansah *et al*. 2008) or an oxidative stress reporter, GstD-GFP (Sykiotis and Bohmann 2008). Neither assay revealed an elevation of oxygen species in iECs but did in discs that expressed the positive control UAS-PDGF- and VEGF-receptor related (*Pvr*), which is known to elevate oxygen species (Figure. S2a-f’) (Wang *et al*. 2016). Moreover, knocking down *Ask1* did not alter JNK activity indicating that Ask does not mediate activation of JNK in iEC (Figure. S2g-h’). All together, these data suggest that ROS and H_2_O_2_ do not play a major role in activating JNK in iECs.

### A Src42A-Shark-Slpr pathway activates JNK activity in iECs

In contrast to the negative results for TNF and ROS pathways, we found that a conserved Src oncogene pathway regulates JNK in iEC (Figure 2a). Src proteins are non-receptor tyrosine kinases that signal through downstream Shark and Slpr proteins to activate JNK in a number of contexts including wound healing, morphogenesis, oncogenesis, and metastasis, functions that are conserved from *Drosophila* to humans (Thomas and Brugge 1997; Gao *et al*. 2004; Rios-Barrera and Riesgo-Escovar 2013). Knockdown of *Src oncogene at 42A (Src42A-i)* (Takahashi *et al*. 1996), the *SH2 ankyrin repeat kinase (Shark-i)* (Ferrante *et al*. 1995), or *slipper (slpr-i)*, a Jun Kinase Kinase Kinase (JNKKK) (Stronach and Perrimon 2002), strongly decreased JNK activity in iECs (Figure 2b-e’’). Overexpression of a negative regulator of Src, UAS-*C-terminal Src kinase* (*Csk*) (Read *et al*. 2004), also strongly inhibited JNK activity in iEC (Figure 2f-f’’). However, knockdown of the only other Src paralog in *Drosophila*, *Src64B (Src64B-i),* (Simon *et al*. 1983) did not alter JNK activity in iECs (Figure. 2g-g’’). These results suggest that a Src42A-Shark-Slpr pathway is the major activator of JNK activity in iEC.

**Figure 2.**
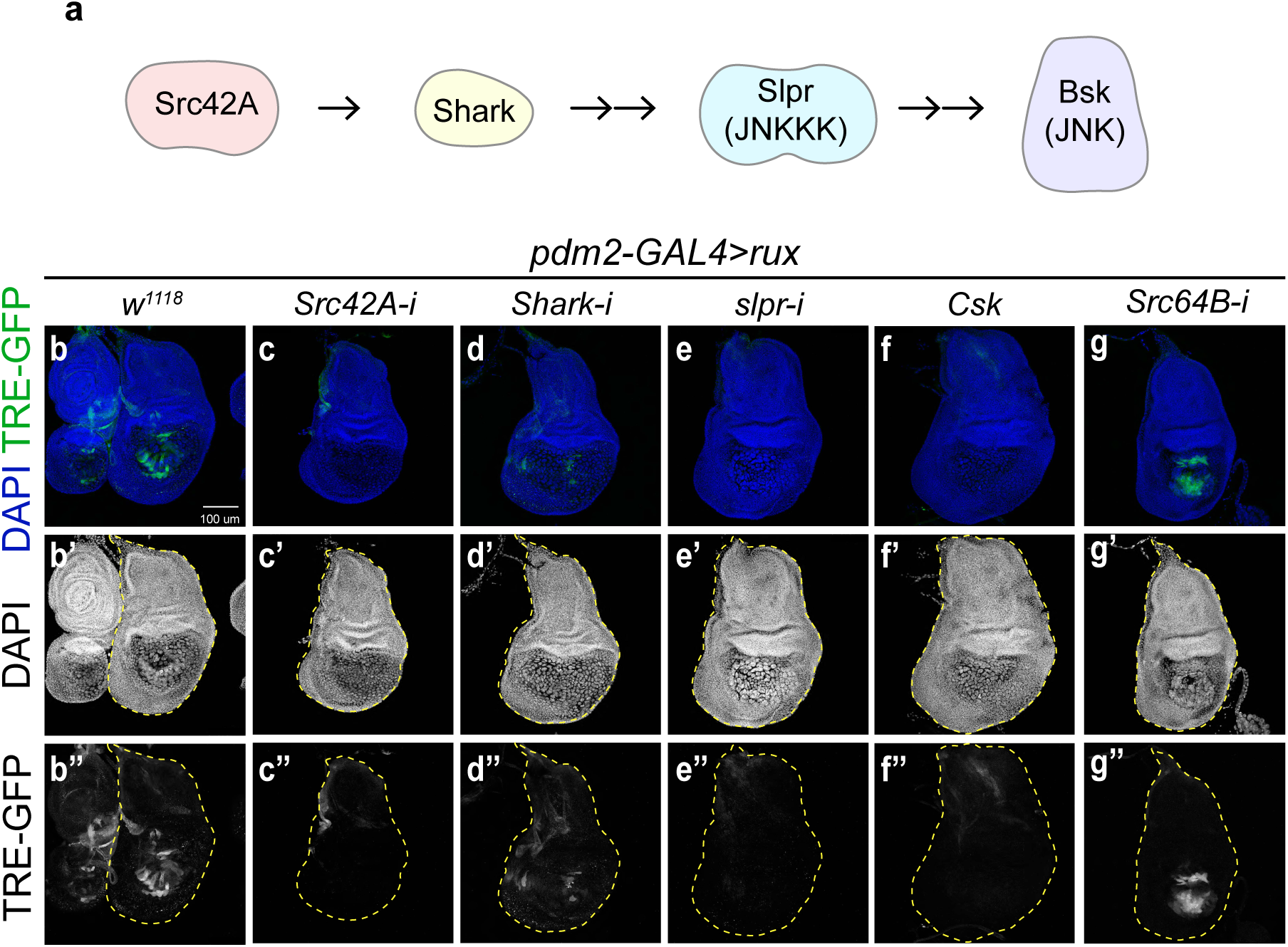
A Src42A-Shark-Slpr pathway activates JNK activity in iECs. (a) Src-JNK signaling pathway in *Drosophila*. (b-g’’) Src42A, Shark, Slpr, but not Src64B, are required to activate JNK in iECs. Confocal images of wing discs with a JNK reporter, TRE-GFP, expressing *UAS-rux* alone (*w^1118^*, b-b’’), or co-expressing UAS-*Src42A^RNAi^* (*Src42A-i*, c-c’’), UAS-*Shark^RNAi^* (*Shark-i*, d-d’’), UAS*-slpr^RNAi^* (*slpr-i*, e-e’’), UAS-*Csk* (*Csk*, f-f’’), or UAS-*Src64B^RNAi^*(*Src64B-i*, g-g’’). Scale bar: 100 µm.

### Src42A-Shark-Slpr induce a senescence-like growth arrest of iEC

We have previously reported that the hypertrophic cell growth of iECs is limited by a JNK-dependent senescence-like arrest (Huang *et al*. 2024b). Given our evidence that the Src42A-Shark-Slpr pathway activates JNK in iECs, we therefore asked if it enforces a senescent growth arrest. Consistent with our previous results, arrested iECs had elevated senescence associated beta-galactosidase (SA-β-gal) activity, which was inhibited by expressing a dominant negative JNK (*bsk^DN^*) (Figure 3a-c”) (Valieva *et al*. 2022). SA-β-gal activity was inhibited to a similar extent by single knockdowns of *Src42A*, *Shark* or *slpr* (Figure 3d-f”). We also measured the ploidy of iECs to assess effects on genome duplications during endocycle progression. Consistent with our previous results, overexpression of UAS-*rux* alone resulted in a population of cells that arrested after different numbers of endocycles and with different terminal ploidies (Figure 3g) (Huang *et al*. 2024b). Co-expression of UAS-*bsk^DN^*, UAS-*Src42A^RNAi^ (Src42A-i)*, UAS-*Shark^RNAi^ (Shark-i)*, or UAS-*slpr^RNAi^* (*slpr-i*) all resulted in iEC populations that had a significantly higher average DNA content than *rux* alone, indicating that they had undergone more endocycles (Figure 3g). These results suggest that a Src42A-Shark-Slpr-JNK pathway induces a senescence-like growth arrest of unscheduled endocycling wing disc cells.

**Figure 3.**
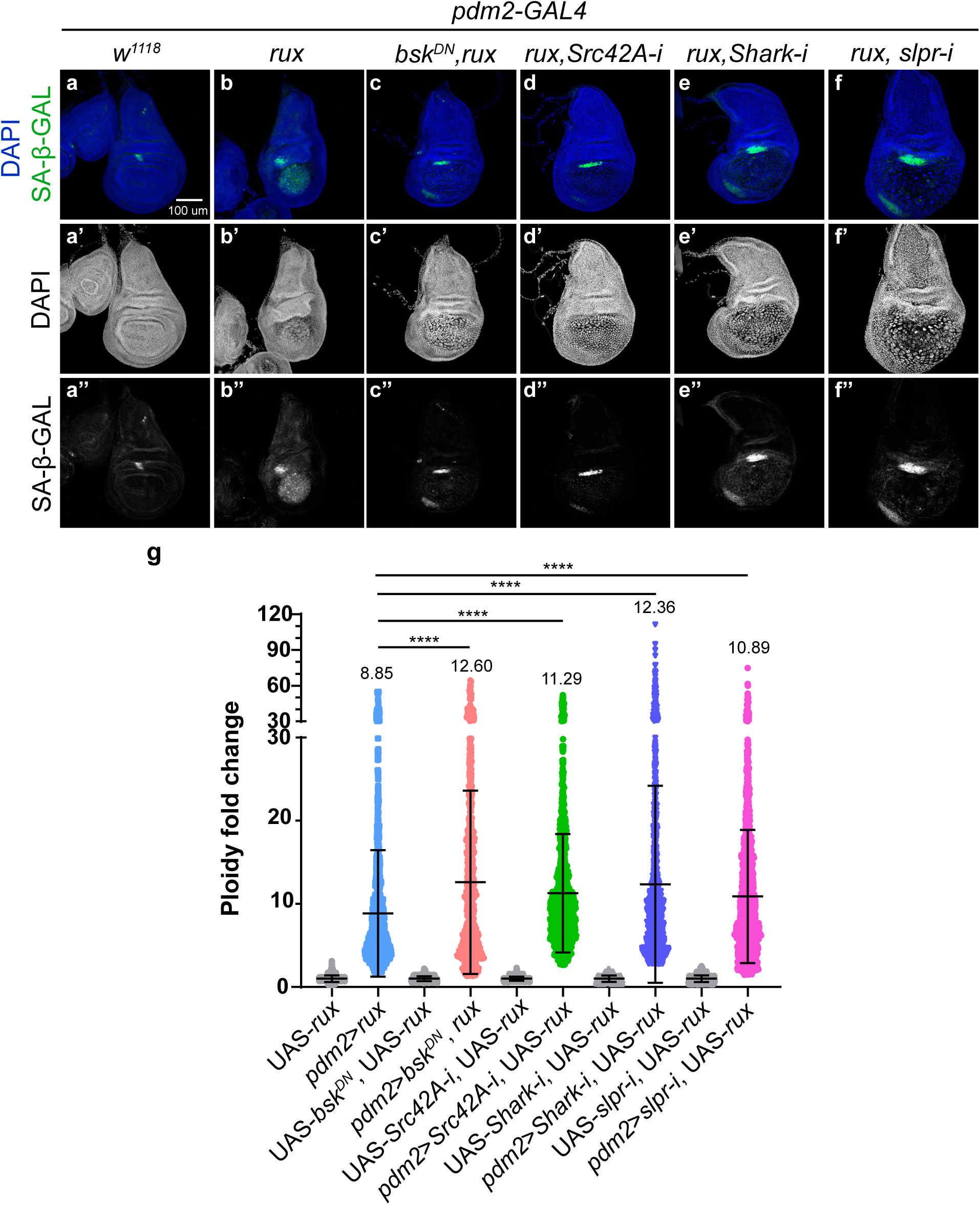
Src42A-Shark-Slpr pathway induces a JNK-dependent senescence-like growth arrest of iECs. (a-f’’) Senescence-associated-β-galactosidase (SA-β-GAL) activity was assayed in the wing disc with *pdm2*-GAL4 alone (*w^1118^*, a-a’’), UAS-*rux* (*rux*, b-b’’), UAS-*rux* and UAS-*bsk^DN^*(*bsk^DN^, rux*, c-c’’), UAS-*rux* and UAS-*Src42A^RNAi^* (*rux, Src42A-i*, d-d’’), UAS-*rux* and UAS-*Shark^RNAi^* (*rux, Shark-i*, e-e’’), or UAS-*rux* and UAS-*slpr^RNAi^* (*rux, slpr-i*, f-f’’). Scale bar: 100 µm. (g) Fold change in DNA content (DAPI fluorescence) of individual iECs (color dots) relative to diploid cells (grey dots) from the same wing disc. **** p<0.01.

### Unscheduled endocycling induces a tissue and organismal wounding-like response

Our results for iEC undergrowth in wing discs suggested similarities to wounding and regeneration responses that are conserved from *Drosophila* to humans. These similarities include activation of a Src-Shark-Slpr-JNK pathway and a senescent-like arrest of a subset of cells at wound sites (Gao *et al*. 2004; Jaiswal *et al*. 2023). In mammals, senescent cells at wound sites promote cell proliferation and wound closure non-autonomously by secreting cytokines including IL-6, a ligand of the JAK/STAT signaling pathway (Demaria *et al*. 2014; Ritschka *et al*. 2017; Andrade *et al*. 2022). Similarly, wound healing and regeneration in *Drosophila* is promoted by expression of the JAK/STAT ligand unpaired 3 (Upd3) (Katsuyama *et al*. 2015; Cosolo *et al*. 2019; Joy *et al*. 2021; Worley and Hariharan 2022; Floc’hlay *et al*. 2023; Jaiswal *et al*. 2023). To further explore similarities to wound healing and regeneration, we determined whether iEC undergrowth activates the JAK/STAT pathway. Induction of unscheduled endocycles with *pdm2*-GAL4>*rux* increased expression of a transcriptional reporter for the Upd3 ligand (*upd3*-lacZ) and a reporter for cells receiving the JAK/STAT signal (STAT-GFP) (Figure S3a-b”, d-e’’). Interestingly, *upd3* expression was highest in the central pouch region whereas STAT92E activity was highest in the hinge cells surrounding the pouch, suggesting that the Upd3 signal produced by the central pouch cells is received by the hinge cells (Figure S3).

Another known response of wing discs to wounding is production of insulin-like peptide 8 (Ilp8), which locally regulates cell proliferation in the wounded tissue and systemically enforces a developmental delay (Colombani *et al*. 2012). To evaluate the timing of the wounding response to unscheduled endocycles, we induced endocycles for different amounts of time using the temperature sensitive GAL4 repressor *tub*-*GAL80^ts^* with *pdm2*-*GAL4*>*rux*, and monitored expression of an Ilp8-GFP transcriptional reporter and the JNK reporter TRE-GFP (Figure 4a) (Garelli *et al*. 2012). The expressions of Ilp8-GFP and TRE-GFP were both first detectable with similar timing, at 48 hours after endocycle induction, with increased expression of both reporters at 72 and 96 hours after endocycle induction (Figure 4b-k’). This timing is consistent with our previous evidence that populations of iEC initially endocycle and grow by hypertrophy for approximately two days before some begin to activate JNK and engage a senescent-like arrest (Huang *et al*. 2024b). The co-temporal JNK activity and Ilp8 expression is consistent with previous evidence that JNK induces Ilp8 expression after wounding (Colombani *et al*. 2012). Co-expression of the dominant negative JNK, *UAS-bsk^DN^*, indicated Ilp8 expression is also dependent on JNK activity in iEC (Figure 4l-m’).

**Figure 4.**
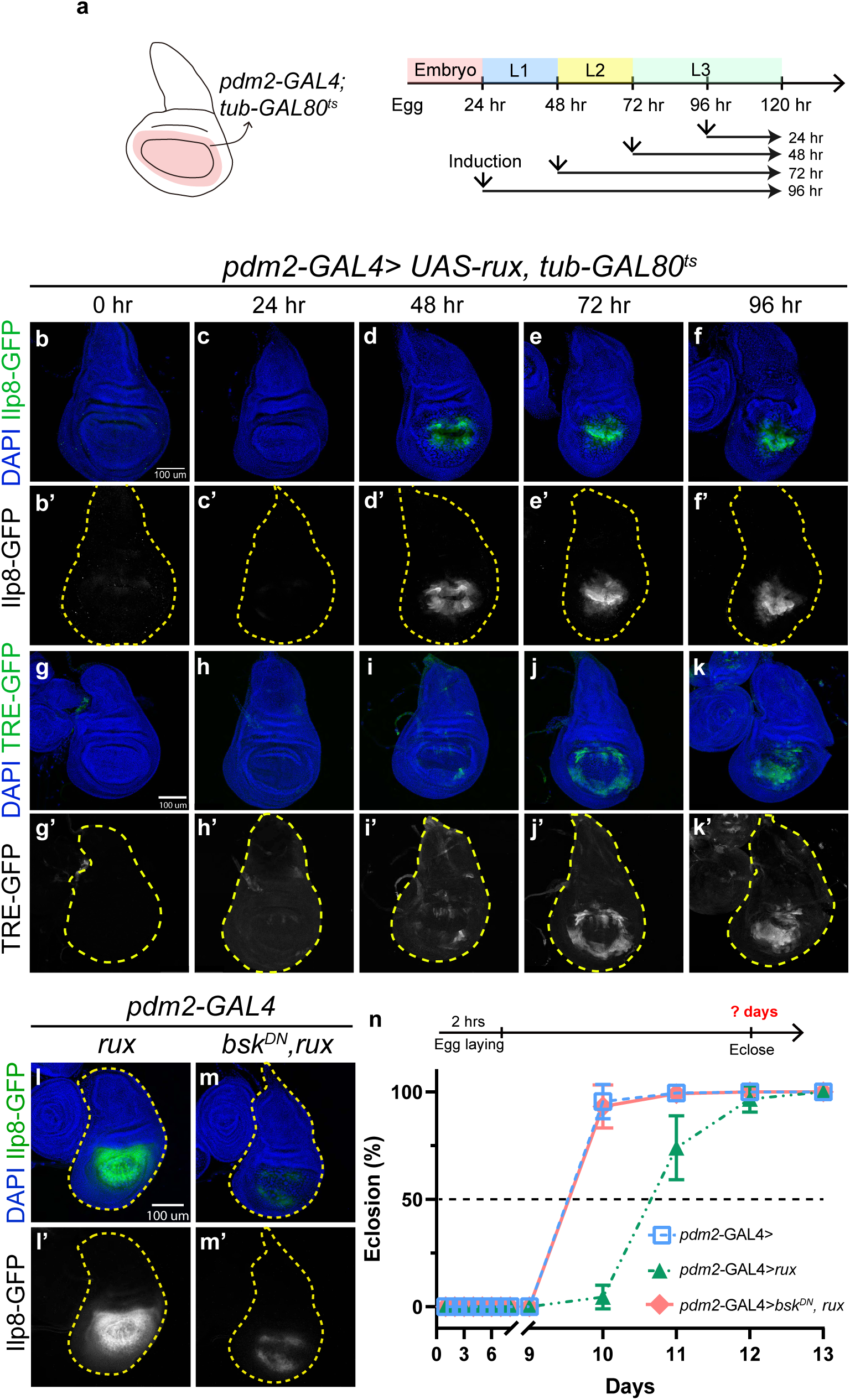
Activation of JNK in iECs induces Ilp8 expression and a developmental delay. (a) Experimental strategy to induce iEC growth for different amounts of time before analysis. (a, left). *Pdm2*-GAL4 was used to induce iECs in pouch and inner hinge (pink). (a, right) *pdm2*-GAL4 activity was inhibited by raising larvae at 18 °C, the GAL80^ts^ permissive temperature, and then induced at different times by shifting from 18 °C to 29 °C, followed by wing disc dissection during late wandering 3^rd^ instar (L3). (b-k’) Ilp8-GFP (b-f’) and JNK reporter, TRE-GFP, expression (g-k’) were analyzed after different durations of iEC growth: 0 hour (b-b’, g-g’), 24 hour (c-c’, h-h’), 48 hour (d-d’, i-i’), 72 hour (e-e’, j-j’), or 96 hour (f-f’, k-k’). Scale bar: 100 µm. (l-m’) Ilp8 expression is JNK-dependent in iECs. Ilp8-GFP expression in the wing disc with UAS-*rux* alone (*rux*, l-l’) or UAS-*rux* and UAS-*bsk^DN^*(*bsk^DN^, rux*, m-m’). Scale bar: 100 µm. (n) Wing disc iECs induce a JNK-dependent developmental delay. Flies with *pdm2*-GAL4 only (*pdm2-* GAL4>, blue), with UAS-*rux* (*pdm2-*GAL4*>rux*, green), or with UAS-*rux* and UAS-*bsk^DN^*, (*pdm2-* GAL4*>bsk^DN^, rux*, red) were allowed to lay eggs for 2 hours and the time to eclosion as adults was quantified every 24 hours.

It is known that release of Ilp8 from wounds acts as a systemic signal to inhibit ecdysone production and enforce a developmental delay of metamorphosis (Garelli *et al*. 2012). To further explore similarities between iEC undergrowth and wounding, therefore, we measured the effect of iECs on the timing of eclosion of adults from the pupal case. Induction of endocycles with *pdm2*-GAL4>*rux* resulted in a significant developmental delay by approximately one day (Figure 4n). This developmental delay was completely abrogated by co-expression of *UAS-bsk^DN^*, which is consistent with our evidence that Ilp8 expression in iECs depends on JNK activity (Figure 4l-m’).

It has been reported that *upd3* expression can also be activated by JNK and can act as a systemic signal that delays development (Katsuyama *et al*. 2015; Romao *et al*. 2021). In wing disc iECs, however, *upd3* expression and its activation of the JAK/STAT reporter were not reduced when JNK activity was inhibited with *bsk^DN^*, indicating that *upd3* expression and JAK/STAT signaling is not dependent on JNK in iECs (Figure S3c-c’’, f-f’’). Moreover, given that inhibiting JNK reverted the developmental delay, this result also suggests that Upd3 is not sufficient to delay development in response to iEC wing disc undergrowth. These results indicate that the responses to iEC undergrowth have many similarities, but also differences, to previously defined wounding and regeneration responses (see discussion).

### Unscheduled endocycling and JNK activation affect wing disc morphology

The wound healing response entails extensive remodeling of cell and tissue morphology. We therefore asked how unscheduled endocycling cells affect tissue morphology by analyzing the three-dimensional structure of the wing disc. The epithelial DP layer forms three folds in specific locations: the hinge-notum (H/N) fold, the hinge-hinge (H/H) fold, and the hinge-pouch (H/P) fold (Figure 5a). We analyzed wing disc morphology by confocal imaging in the x-y and y-z axes (Figure 5a). Confocal imaging of *pdm2*-*GAL4*>*RFP* alone confirmed that *pdm2-GAL4* is expressed in the wing pouch and in part of the inner wing hinge that includes the H/P fold (Figure 5b-b’, f-f’, video 1) (Butler *et al*. 2003; Jory *et al*. 2012; Loker and Mann 2022). In the posterior wing disc compartment, the *pdm2*-*GAL4* expression domain extends into the cuboidal cells at edge of the wing disc (Figure 5b-b’, video 1). Induction of endocycles with *pdm2*-GAL4>*rux* resulted in undergrowth of this RFP+ region, but the three distinct wing disc folds still formed (Figure 5c-c’, g-g’, video 2). The pouch cells at the H/P border also formed a distinct circular structure in the x-y axis (Figure 5c-c’). Imaging in the x-y and y-z axes indicated that this circular morphology is the result of circumferential folding and centripetal extension of the hinge tissue under the basal side of the pouch epithelium (Figure 5c-c’, g-g’, video 2). This extension was associated with an increased number of RFP-diploid hinge cells outside of the *pdm2*-GAL4 domain, consistent with our previous findings that iEC undergrowth stimulates compensatory proliferation of neighboring cells (Figure 5g-g’, video 2) (Huang *et al*. 2024b).

**Figure 5.**
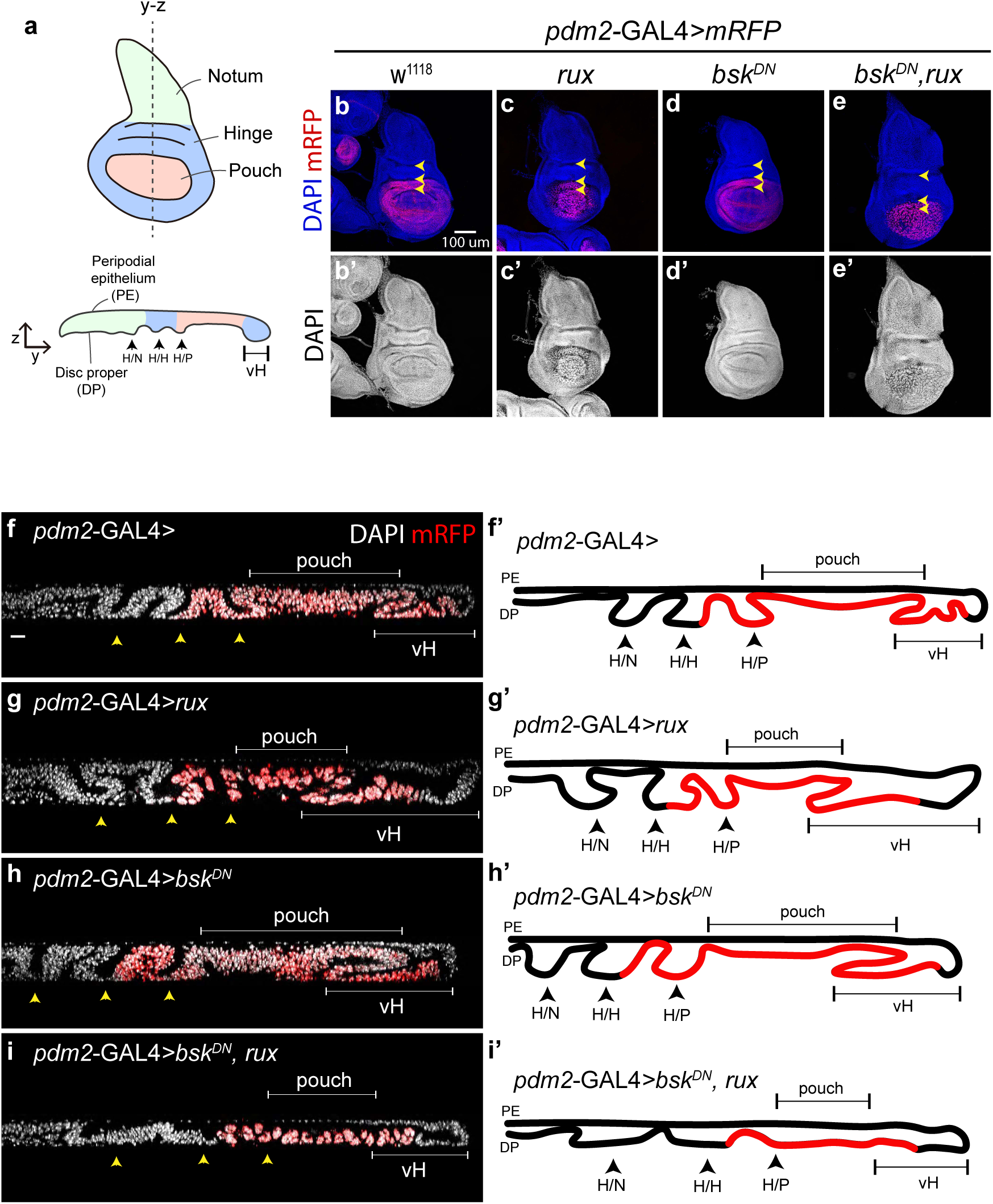
Unscheduled endocycling and JNK activation affect wing disc morphology. (a) Morphology of *Drosophila* wing disc. The top drawing is a wing disc in the x-y plane. Wing discs have three distinct epithelial folds between notum, hinge, and pouch: notum/hinge (N/H) fold, hinge/hinge (H/H) fold, and hinge/pouch (H/P) fold. vH: ventral hinge. The bottom drawing is a transverse slice through the wing disc in the y-z plane, which is highlighted as a dashed line in the top drawing. The peripodial epithelium (PE) is on top and the disc proper epithelium (DP) is on the bottom with DP apical side of cell up. The three folds and ventral hinge are indicated as in the top drawing. (b-e) Wing discs imaged in the x-y plane with *pdm2*-GAL4 only (*w^1118^*, b-b’), UAS-*rux* (*rux*, d-d’), UAS-*bsk^DN^*, (*bsk^DN^*, d-d’), or UAS-*rux* and *bsk^DN^*, (*bsk^DN^*, *rux*, e-e’). Cells expressing *pdm2*-GAL4 are marked by UAS-*mRFP* expression. Arrow heads mark the three folds shown in (a). Scale bar: 100 µm. (f-i’) Transverse section across the wing disc in the y-z plane with interpretation drawings of micrographs. *pdm2*-GAL4 expressing cells are marked by UAS-*mRFP* and three folding structures: H/N, H/H, and H/P are indicated by arrow heads. Scale bar: 20 µm.

It is known that JNK mediates reorganization of tissue morphology during wound healing and development. We therefore asked whether JNK activation contributes to the altered morphology of wing discs with iECs. Expression of *pdm2-*GAL4*>bsk^DN^* in diploid control discs did not alter the formation of the H/P, H/H, H/N folds (Figure 5d-d’, h-h’, video 3). In contrast, inhibition of JNK kinase activity in iEC discs (*pdm2*-GAL4*> bsk^DN^,rux*) greatly reduced fold formation resulting in a flatter DP epithelium and reduced expansion of the diploid hinge region (Figure 5e-e’, i-i’, video 4). These results suggest that JNK activity is required for enhanced epithelial folding and hinge tissue extension in iEC wing discs.

### A Src-JNK pathway induces actomyosin in H/P fold iECs and centripetal extension of hinge tissue under the central pouch

To further define the relationship between JNK activity and the effect of iECs on wing disc morphology, we analyzed the spatial and temporal correlation between JNK activity and wing disc folding. We used the temperature sensitive GAL4 inhibitor, GAL80^ts^, with *pdm2*-GAL4>*rux* to induce iEC growth for different lengths of time and then analyzed the expression of the JNK reporter, TRE-GFP, in the x-y and y-z axes. Similar to our results in Figure 3, TRE-GFP was weakly and variably expressed in a ring of hinge cells beginning at 48 hrs, with expression increasing at 72 and 96 hours (Figure 6a-e”). Imaging in the y-z axes showed that this ring of TRE-GFP positive cells formed the leading edge of the hinge tissue that extends centripetally under the pouch (Figure 6f-j). At 96 hours, these leading-edge TRE-GFP+ H/P cells extended even farther underneath the pouch (Figure 6e-e”, j). These results show that the cells with highest JNK activity form the leading edge of a progressively elongating polyploid hinge tissue.

**Figure 6.**
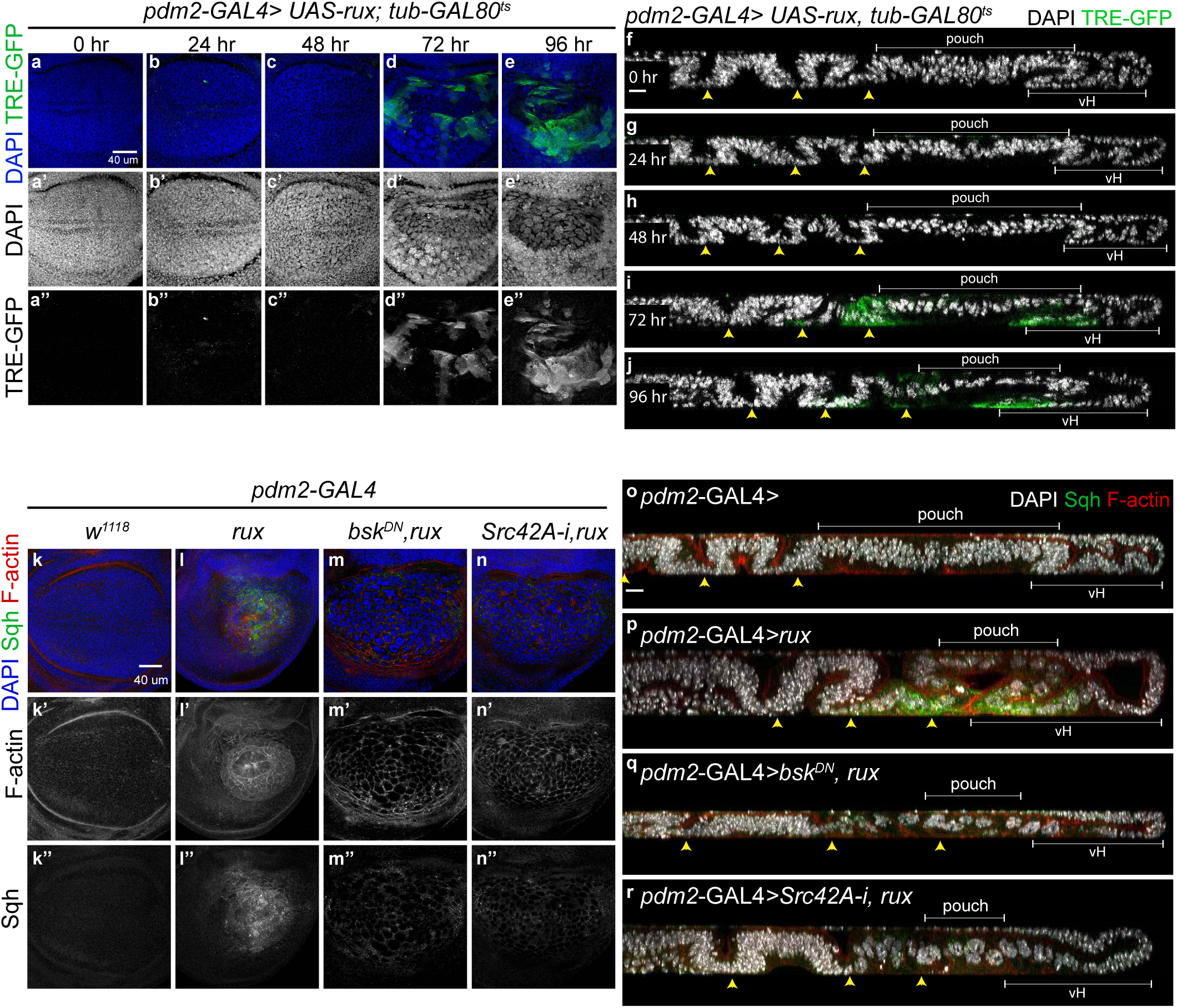
The Src-JNK pathway induces actomyosin at leading edge of hinge folds. (a-j) Temporal and spatial activity of JNK in iECs. 3rd instar wing discs with *pdm2*-GAL4, UAS-*rux*, *tub*-GAL80^ts^ were incubated in 29 °C for different periods before analyzing TRE-GFP expression. The images are shown in the x-y plane (a-e’’) and in the y-z plane (f-j). Scale bar: 40 µm for (a-e’’) and 20 µm for (f-j). (k-r) Actomyosin expression in iECs. Larvae were raised in 25 °C and wing discs were collected at late 3rd instar with *pdm2*-GAL4 alone (*w^1118^*, k-k’’, o), UAS-*rux* (*rux*, l-l’’, p), UAS-*rux* and UAS-*bsk^DN^*(*bsk^DN^, rux*, m-m’’, q), or UAS-*rux* and UAS-*Src42A^RNAi^* (*rux, Src42A-i*, n-n’’, r). F-actin was labeled with fluorescent phalloidin (red) and non-muscle type 2 myosin (Sqh, green) was detected by using a reporter strain (Sqh-GFP). Scale bar: 40 µm for (k-n’’) and 20 µm for (o-r).

It is known that the Src-JNK pathway regulates actomyosin expression to induce cell motility and tissue reorganization in a variety of biological contexts (Thomas and Brugge 1997; Vidal *et al*. 2007). To further investigate how JNK induces extension of hinge tissue, therefore, we examined the expression of a transcriptional reporter for the non-muscle myosin light chain, *spaghetti squash (sqh)* (Karess *et al*. 1991), and the formation of filamentous actin (F-actin) by labeling with fluorescent phalloidin (Wulf *et al*. 1979). Induction of unscheduled endocycles increased *sqh* expression and F-actin formation, which was greatest in the extending H/P fold cells that had high JNK activity (Figure 6i-j, k-l’’, p). Imaging in the x-y axis revealed that F-actin was organized among adjacent cells to form a larger, distinct ring structure, reminiscent of the previously described supracellular actomyosin cables that facilitate *Drosophila* embryonic dorsal closure and wound healing (Figure 6k-l’’) (Jacinto *et al*. 2002; Omelchenko *et al*. 2003; Hayes and Solon 2017). Imaging in the y-z axis indicated that this actomyosin ring structure is being formed by the centripetal extension of hinge tissue (Figure 6p). In some cases, the hinge cells from different circumferential positions joined at the center similar to tissue suturing during wound closure and the dorsal closure of the *Drosophila* embryo (Figure 6p) (Huang *et al*. 2024a). Expression of *Sr42A^RNAi^* or *bsk^DN^* resulted in lower *sqh* expression and F-actin formation indicating that they are dependent on activation of the Src-JNK pathway (Figure 6m-n’’, q-r).

Inhibition of Src42A and JNK activity also disrupted the circular arrangement of cells and F-actin in the x-y axis and the formation of the hinge underfold seen in the y-z axis (Figure 6q-r). These results suggest that the Src-JNK-actomyosin pathway in polyploid H/P fold cells induces a cytoskeletal and tissue reorganization that is similar to that of wound closure.

### Cell adhesion molecules are upregulated in central pouch iECs

The results indicated that unscheduled endocycles activate a Src-JNK-actomyosin pathway that elicits an extensive reorganization of tissue morphology. To explore this further, we examined cell adhesion molecules that play important roles in tissue dynamics during morphogenesis and wound healing (Bulgakova *et al*. 2012; Besen-Mcnally *et al*. 2021). We first examined expression of integrins which transduce cell and tissue forces by mediating cell-extra cellular matrix (ECM) and cell-cell interactions (Plotnikov and Waterman 2013). We used a reporter strain for the *myospheroid (mys)* gene, the beta subunit of integrin, which expressed a Mys-GFP fusion protein encoded by the endogenous *mys* locus (Klapholz *et al*. 2015). Mys-GFP expression was increased in the iEC of the central pouch and formed distinct puncta, consistent with previous evidence that integrin proteins coalesce into focal adhesions to mediate cell-ECM interactions and transduce tissue forces (Figure 7a-b’’, arrow heads) (Katsumi *et al*. 2005; Pereira *et al*. 2011). While Mys-GFP expression and localization was clearly altered in the central pouch cells, its expression was not detectably increased in the surrounding H/P fold iECs (Figure 7b-b”). We next examined the expression of *four-jointed (fj)*, a trans-membrane kinase that phosphorylates and regulates the noncanonical cadherin proteins Fat (Ft) and Dachsous (Ds) (Ishikawa *et al*. 2008). These proteins are part of non-canonical adherens junctions that mediate cell-cell interactions for planar cell polarity, growth, and regeneration. The transcriptional reporter for *fj* (*fj-lacZ*) was upregulated in the central pouch cells, but not the adjacent H/P fold iECs, identical to the expression pattern of *mys* (Figure 7d-e’’) (Brodsky and Steller 1996).

**Figure 7.**
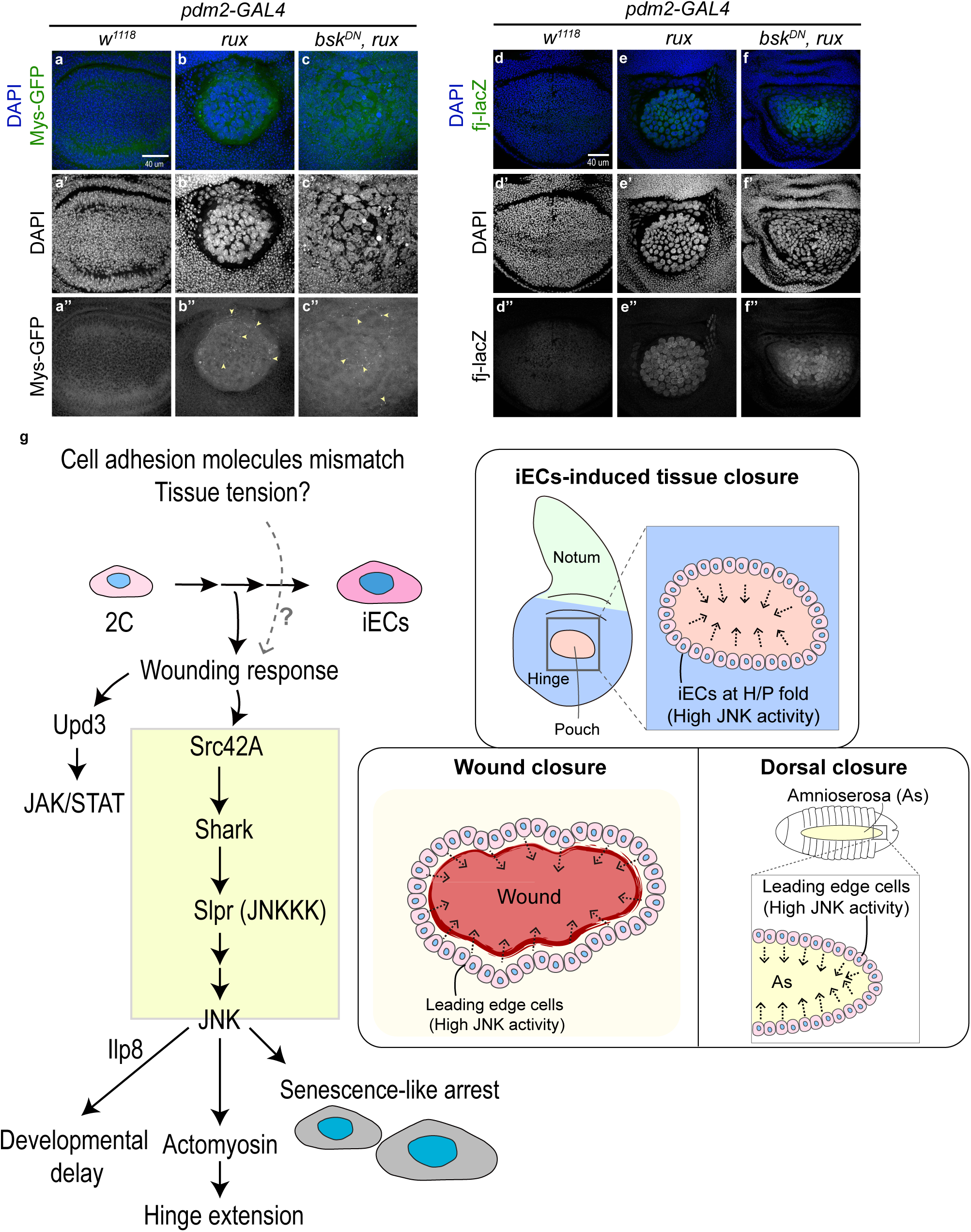
Integrin and Fj expression are upregulated in unscheduled endocycling pouch cells. (a-f’’) Confocal images of wing disc with *pdm2*-GAL4 only (*w^1118^*, a-a’’, d-d’’), UAS-*rux* (*rux*, b-b’’, e-e’’), or UAS-*rux* and UAS-*bsk^DN^* (*bsk^DN^*, *rux*, c-c’’, f-f’’). Expression of *myospheroid* (*mys*) expression was detected with the mys-GFP reporter (a-c’’) and *four-jointed* (*fj*) by fj-GFP reporter (d-f’’). Focal adhesions are indicated by yellow arrow heads in b”, c”. Scale bar: 40 µm. (g) Summary and model for tissue response to iECs through a wound-like response. This response entails activation of a Src-JNK pathway, which enforces iEC senescent arrest, produces wounding signals with systemic developmental delay, and reorganizes actomyosin and tissue morphology. The expression pattern and epithelial morphology changes in iEC discs are similar to those induced by the JNK pathway in wound healing and *Drosophila* embryonic dorsal closure. See discussion for details.

The results indicated that *mys* and *fj* were upregulated in the central pouch, but not the H/P fold iECs that had high actomyosin expression and that participate in extensive tissue remodeling. This result was unexpected given that integrins are known to link the actin cytoskeleton with the ECM to mediate cell migration, morphogenesis, and wound healing (Katsumi *et al*. 2005; Goodwin *et al*. 2016). Moreover, during those and other processes, JNK has been shown to induce integrin expression, yet integrin was not detectably increased in the H/P fold cells with the highest JNK expression (Karkali *et al*. 2023). To address this question further, we inhibited JNK activity in iEC with *bsk^DN^* and found that it did not alter the expression of Mys-GFP or fj-lacZ in the central pouch cells, indicating that induction of *mys* and *fj* expression in these iECs is not dependent on JNK. (Figure 7c-c’’, f-f’’). These results indicate that the effects of iECs on wing disc morphology are associated with elevated *Mys* and *Fj*, two proteins known to have roles in tissue morphogenesis and regeneration.

## Discussion

Responses to wounding and impaired growth are essential for normal tissue development and homeostasis. One challenge to tissue growth is an unscheduled switch to polyploid endocycles, but how tissues respond to this challenge is ill-defined. Here, using the *Drosophila* wing disc as model, we found that a Src42A-JNK pathway is activated in induced endocycling cells (iECs) and promotes their senescent growth arrest. Activation of this Src-JNK pathway also altered the actomyosin cytoskeleton and three-dimensional epithelial morphology, similar to the known functions of this pathway in morphogenesis, cell migration, and wound healing (Figure 7g). Also similar to wounding, iECs expressed secreted proteins of the JAK/STAT and insulin-like peptide pathways. These results suggest that unscheduled endocycles induce a wounding response, but with some key differences. Among these differences are that growth-arrested iECs are resistant to cell death and not eliminated from the tissue, thereby acting as a constant source of wounding signals in a futile attempt to regenerate. Overall, our results are broadly relevant to understanding the beneficial and detrimental effects of polyploid cells at wounds, chronic wounds, and tumors.

### Unscheduled endocycles induce Src-JNK and a modified wounding response

Our study revealed that iECs elicited responses with many similarities to wound healing. These similarities included activation of a Src42A-Shark-Slpr-JNK senescent arrest, expression of actomyosin, and expression of two secreted proteins with functions in the wounding response and regeneration – Ilp8 and Upd3. Ilp8 also induced a developmental delay similar to its previously described systemic effects after imaginal disc wounding (Figure 7g) (Garelli *et al*. 2012; Romao *et al*. 2021). The Upd3 ligand of the JAK/STAT signaling pathway has functions similar to mammalian IL-6. IL-6 is secreted by senescent cells at wounds together with 100’s of other proteins, which is collectively known as the senescent associated secretory phenotype (SASP), an expression profile that we previously showed is mostly conserved to senescent *Drosophila* iECs (Demaria *et al*. 2014; Andrade *et al*. 2022; Huang *et al*. 2024b). It is known that JAK/STAT signaling promotes wound healing in part by inducing the compensatory proliferation of cells to replace missing tissue, a regenerative response that we had previously shown is activated by senescent iECs in wing discs (Jiang *et al*. 2009; Katsuyama *et al*. 2015; Huang *et al*. 2024b). Thus, there are striking parallels between the effects of senescent cells at wounds and senescent unscheduled endocycling cells in growing tissues. In addition, we observed elevated expression of the integrin protein, Mys, and a regulator of non-canonical adherens junctions, Fj, both of which have cell nonautonomous effects in morphogenesis and wound healing (Gorfinkiel *et al*. 2009; Liu *et al*. 2010; Wong *et al*. 2011; Montes and Morata 2017; Li *et al*. 2024). Thus, unscheduled endocycling cells induce a response that is similar to that of wounding.

The pattern of Src-JNK activity and dynamic tissue reorganization of iEC discs was also highly reminiscent of wound healing and dorsal closure of the *Drosophila* embryo. During these other processes, high levels of Src-JNK activity induces supracellular actomyosin cables in a ring of cells that form the leading edge of an inward tissue extension that results in a purse string-like closure of wounds or the dorsal embryo (Figure 7g) (Gao *et al*. 2004; Hayes and Solon 2017; Hunter *et al*. 2018). We observed a similar morphology of F-actin in a ring of hinge cells with high Src-JNK activity at the leading edge of a circular tissue closure in a futile effort to replace the growth-arrested pouch (Figure 7g). Thus, from a three-dimensional tissue perspective, the centripetal extension and closure of hinge tissue is similar to Src-JNK regulated morphogenesis and wound healing.

Also similar to wound healing and dorsal closure, the effect of iECs on tissue architecture was associated with regional differences in levels of JNK activity. We had induced endocycles in wing pouch and part of the surrounding hinge with *pdm*-*GAL4*, but only the hinge iECs at the H/P fold had significant expression of the JNK reporter TRE-GFP, an artificial promoter reporter composed of multiple binding sites for the AP-1 transcription factor complex. While most pouch cells did not have detectable TRE-GFP, the expression of SA-β-gal and Ilp8, as well as senescent growth arrest in these pouch cells were all dependent on Src-JNK. We interpret these data to mean that the artificial TRE-GFP promoter reporter does not detect low levels of JNK activity, and that the pouch iECs have lower activity of the Src-JNK pathway than surrounding hinge cells (Chatterjee and Bohmann 2012). The elevated Src-JNK activity in hinge cells was required for actomyosin expression and extension of hinge cells underneath the pouch iECs. This behavior of hinge cells is consistent with previous evidence that they play an important role in regeneration by being death-resistant and migrating centrally into the pouch region to replace dying and delaminating pouch cells (Herrera *et al*. 2013; Verghese and Su 2016; Sun *et al*. 2020). The elevated expression of the JNK reporter TRE-GFP we observed in hinge raises the possibility that senescent-arrested iEC pouch cells may be emitting signals for regeneration that promote hinge JNK activity. Consistent with this suggestion, a similar ring of TRE-GFP hinge expression was previously reported after induction of pouch cell death (Harris *et al*. 2016). We found that iEC pouch cells had highest expression of the JAK /STAT ligand, upd3, whereas hinge cells had highest activity of the STAT92E reporter for cells receiving the JAK /STAT signal. Given that JAK/STAT signaling is known to play an important role in regeneration, this leads to the hypothesis that JAK/STAT signaling from the pouch may induce regenerative activity of the hinge (Katsuyama *et al*. 2015; Jaiswal *et al*. 2023). Distinct from other studies of disc regeneration, however, senescent-arrested pouch iECs neither die nor delaminate, and there was no evidence of hinge cell migration into the pouch region (Huang *et al*. 2024b). One interpretation is that hinge cells are being induced to replace the growth-arrested pouch iECs but are unable to displace them and instead fold underneath the pouch as they try to migrate centripetally. The persistence of pouch iECs may be related to their increase in focal adhesions that anchor them to the basement membrane. Thus, persistent, growth-arrested unscheduled endocycling cells present unique challenges for tissue growth and regeneration.

Although there were many similarities between the response to unscheduled endocycles and wounding, there were also some key differences. The first was that we did not find evidence for ROS production using two reporters, a common property of wounding, nor was JNK activation dependent on Ask1, the protein that mediates ROS activation of JNK after wounding (Rhee 1999; Tobiume *et al*. 2001; Worley and Hariharan 2022). Another notable difference between wounding and unscheduled endocycles was the timing of JNK activation. After wounding, JNK is activated within hours (Ramet *et al*. 2002; Losick *et al*. 2013), whereas we found that JNK was activated only after two days of induced endocycle growth. This observation suggests that there is a cumulative property of repeated unscheduled endocycles that results in a delayed activation of Src-JNK and a downstream senescent arrest and wound response. It is unclear at present what that property is. One possibility is that the excessive growth and unusual shape of endocycling cells perturbs cell-cell interactions that is perceived as a cell surface mismatch that activates the Src pathway. Indeed, we previously found that iECs have profound effects on cell and epithelial shape in the ovary (Herriage and Calvi 2024). Moreover, undergrowth of iEC may induce a mechanical tissue stress that activates the Src-JNK pathway, a mechanism that activates this pathway in other contexts (Katsumi *et al*. 2005; Pereira *et al*. 2011). Altogether, our study has uncovered many molecular, cellular, and tissue similarities between the response to wounding and iECs, but has also revealed properties that are unique to unscheduled endocycles.

### Polyploid cells of wounds, chronic wounds, and tumors

Our findings have broader medical impact because they suggest that the unscheduled endocycles induced by aging and other stresses have the potential to contribute to tissue dysfunction. Our results also raise a paradox because natural induction of unscheduled polyploid cells at wound sites can be beneficial for wound healing (Losick *et al*. 2013; Ovrebo and Edgar 2018; Kirillova *et al*. 2021). Similar to our findings, polyploid cells at wounds have JNK activity and express integrins and cytoplasmic myosin, which contribute to restoring tissue tension after regeneration (Cao *et al*. 2017; Losick and Duhaime 2021). It has been shown that senescent cells are induced at wound sites in mammals and benefit wound healing by secreting IL-6 and other SASP proteins (Ritschka *et al*. 2017; Andrade *et al*. 2022; Jaiswal *et al*. 2023). This leads us to suggest that large, polyploid, senescent cells at wounds may promote healing in part by resisting cell death and producing large quantities of cytokines. Why then are senescent polyploid cells deleterious for wing disc growth? One reason is that iECs persist and cannot be displaced by proliferating diploid cell neighbors during regeneration. Further, unlike an acute wounding response that turns down JNK and cytokines after healing, these persistent iECs have continuous JNK activity and cytokine production, properties that are shared with the known pathological causes of chronic wounds and hyperinflammation (Galko and Krasnow 2004; Hammouda *et al*. 2020). This model is relevant to effects of endocycling Polyploid Giant Cancer Cells (PGCCs) in humans, which also undergo a senescent arrest, are cell death-resistant, and produce cytokines that promote proliferation of neighboring cells (Faggioli *et al*. 2023; Herriage *et al*. 2024; Liu *et al*. 2024). Thus, persistent iECs may be deleterious by inducing a chronic wounding response in both tissues and tumors.

## Acknowledgements

We thank H. Herriage, P. Rangarajan, and C. Steffensen for their feedback on the manuscript. Thanks to the Bloomington fly community and A-K Classen for helpful discussions. We thank E.A. Bach, T. Kaufman, and the Bloomington *Drosophila* Stock Center for fly strains. Thanks to A. Kun from IU Light Microscopy Imaging Center (LMIC) for imaging advice, and FlyBase for essential information.

## Supplemental Material

**Figure S1.**
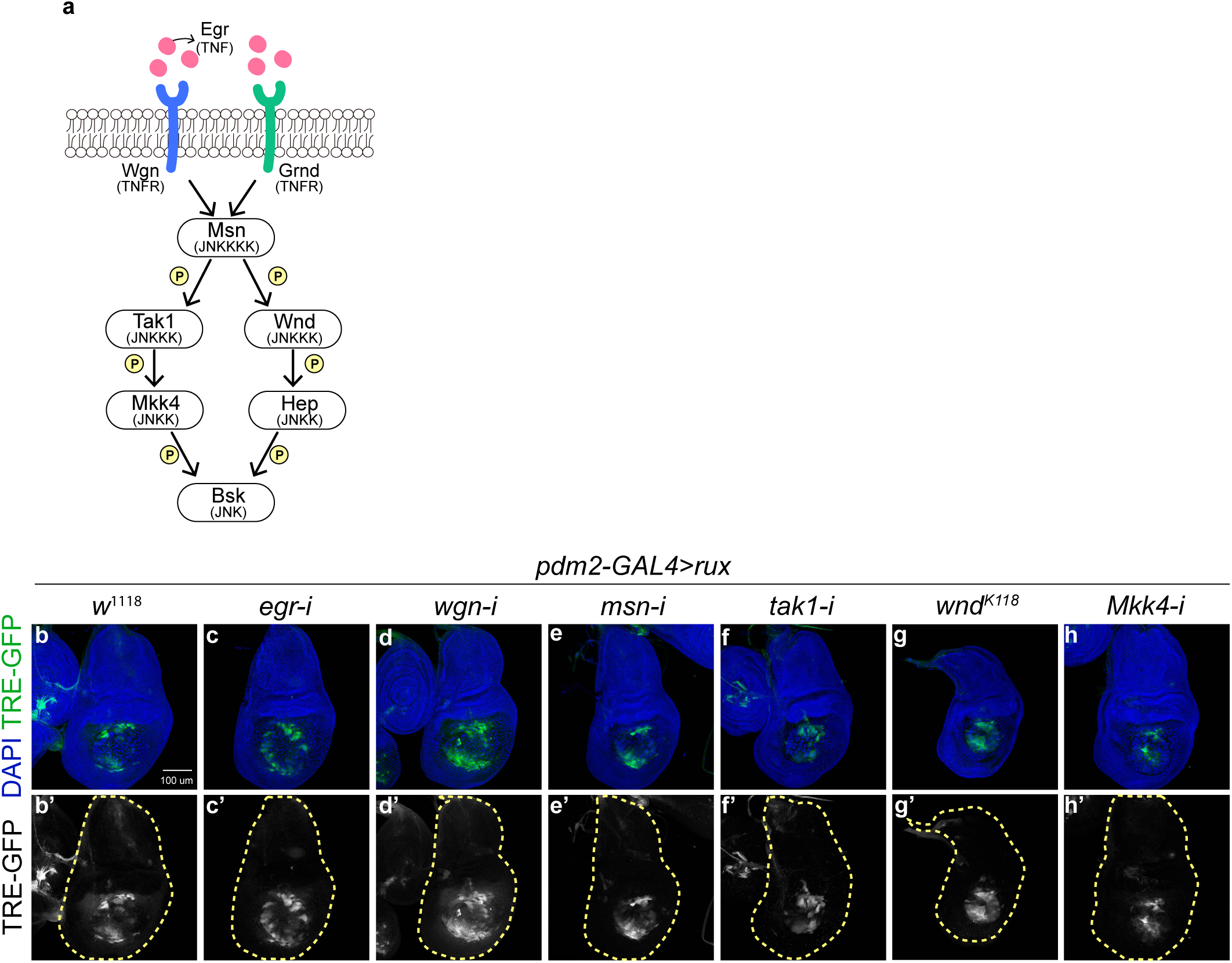
Eiger pathway does not activate JNK in iECs. (a) TNF, *eiger (egr)*, signaling pathway in *Drosophila*. (b-h’) Images of wing discs with a JNK reporter, TRE-GFP, expressing UAS-*rux* (*rux*, b-b’), UAS-*rux* and UAS-*egr^RNAi^* (*rux, egr-i*, c-c’), UAS-*rux* and *wgn^RNAi^* (*rux, wgn-i*, d-d’), UAS-*rux* and UAS-*msn^RNAi^* (*rux, msn-i*, e-e’), UAS-*rux* and UAS-*tak1^RNAi^* (*rux, tak-i*, f-f’), UAS-*rux* and kinase-dead form of UAS-*wnd* (*rux, wnd^k118^*, g-g’), or UAS-*rux* and UAS-*Mkk4^RNAi^*(*rux, Mkk4-i*, h-h’). Scale bar: 100 µm.

**Figure S2.**
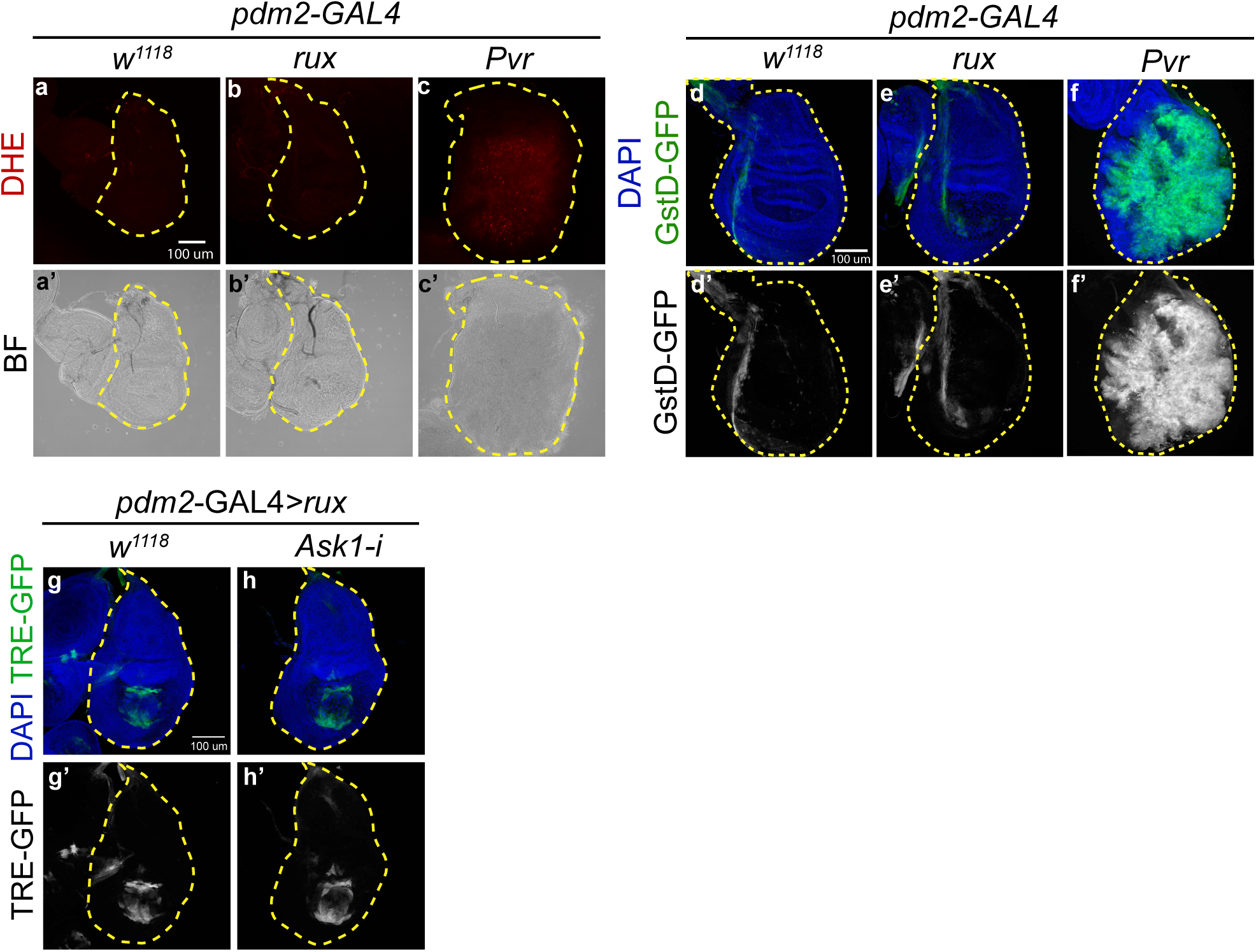
ROS and H_2_O_2_ are not elevated nor activate JNK in iECs. (a-d’) ROS and H_2_O_2_ levels were assayed by dihydroethidium (DHE) staining (a-c’) or by the reporter, GstD-GFP, (d-f’) in wing disc with *pdm2*-GAL4 only (*w*^1118^, a-a’, d-d’), *pdm2*-GAL4 with UAS-*rux* (*rux*, b-b’, e-e’), or *pdm2*-GAL4 with positive control UAS-*Pvr* (*Pvr*, c-c’, f-f’) Scale bar: 100 µm. BF: bright field. Scale bar: 100 µm. (g-h’) Confocal images of wing disc with a JNK reporter, TRE-GFP, expressing UAS-*rux* (*rux*, g-g’), or UAS-*rux* and UAS-*Ask1^RNAi^* (*rux, Ask1*-i, h-h’). Scale bar: 100 µm.

**Figure S3.**
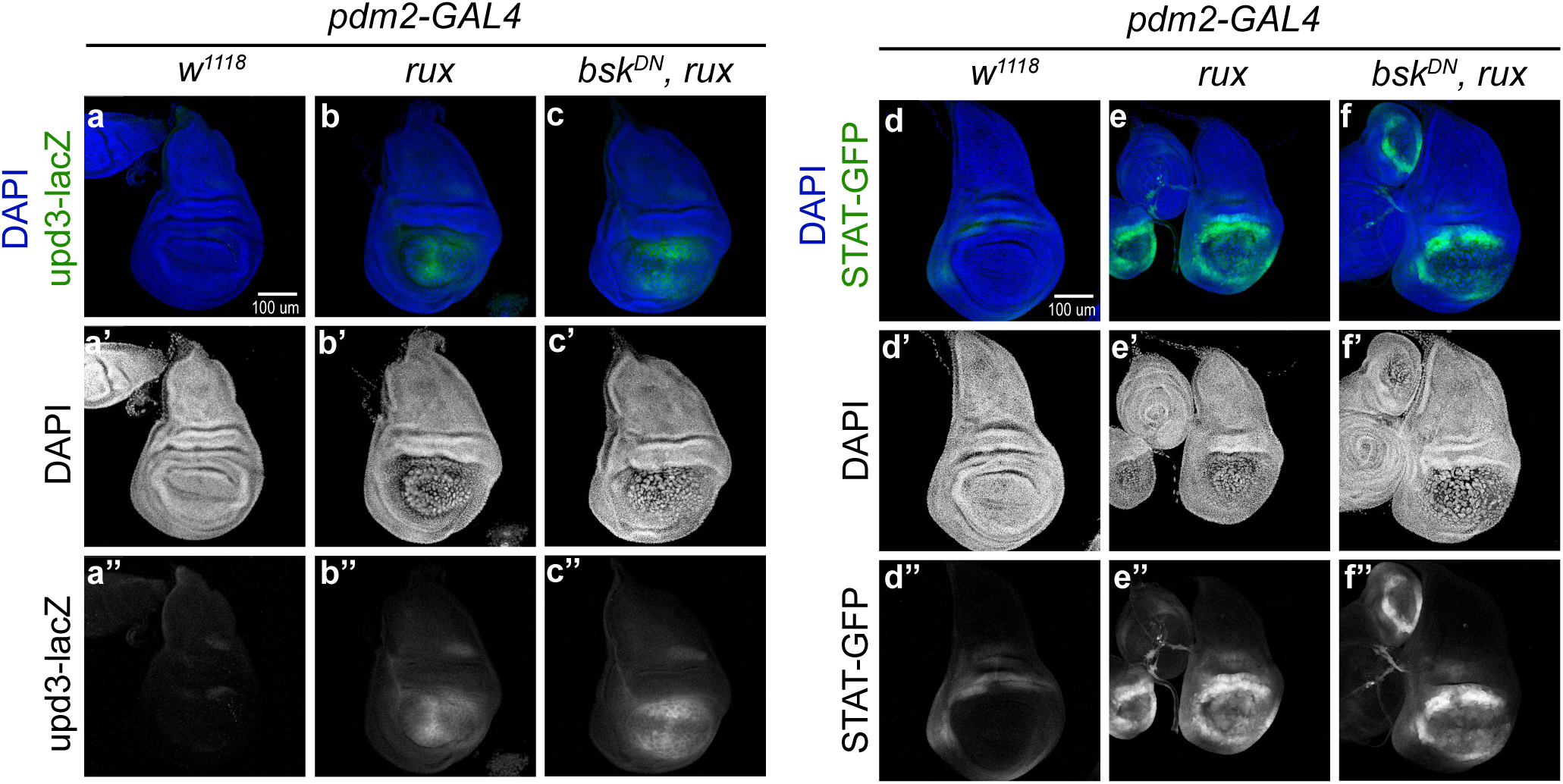
Upd3 and JAK/STAT pathway are upregulated in iECs but not dependent on JNK. Expression of the ligand of JAK/STAT pathway, *upd3*, and the activity of JAK/STAT pathway are detected by reporter strains, *upd3*-*lacZ* (a-c’’) and STAT-GFP (d-f’’), in the wing discs expressing *pdm2*-GAL4 alone (*w^1118^*, a-a’’, d-d’’), UAS-*rux* (*rux*, b-b’’, e-e’’), or UAS-*rux* and UAS-*bsk^DN^* (*bsk^DN^, rux*, c-c’’, f-f’’). Scale bar: 100 µm.

**Table S1:**
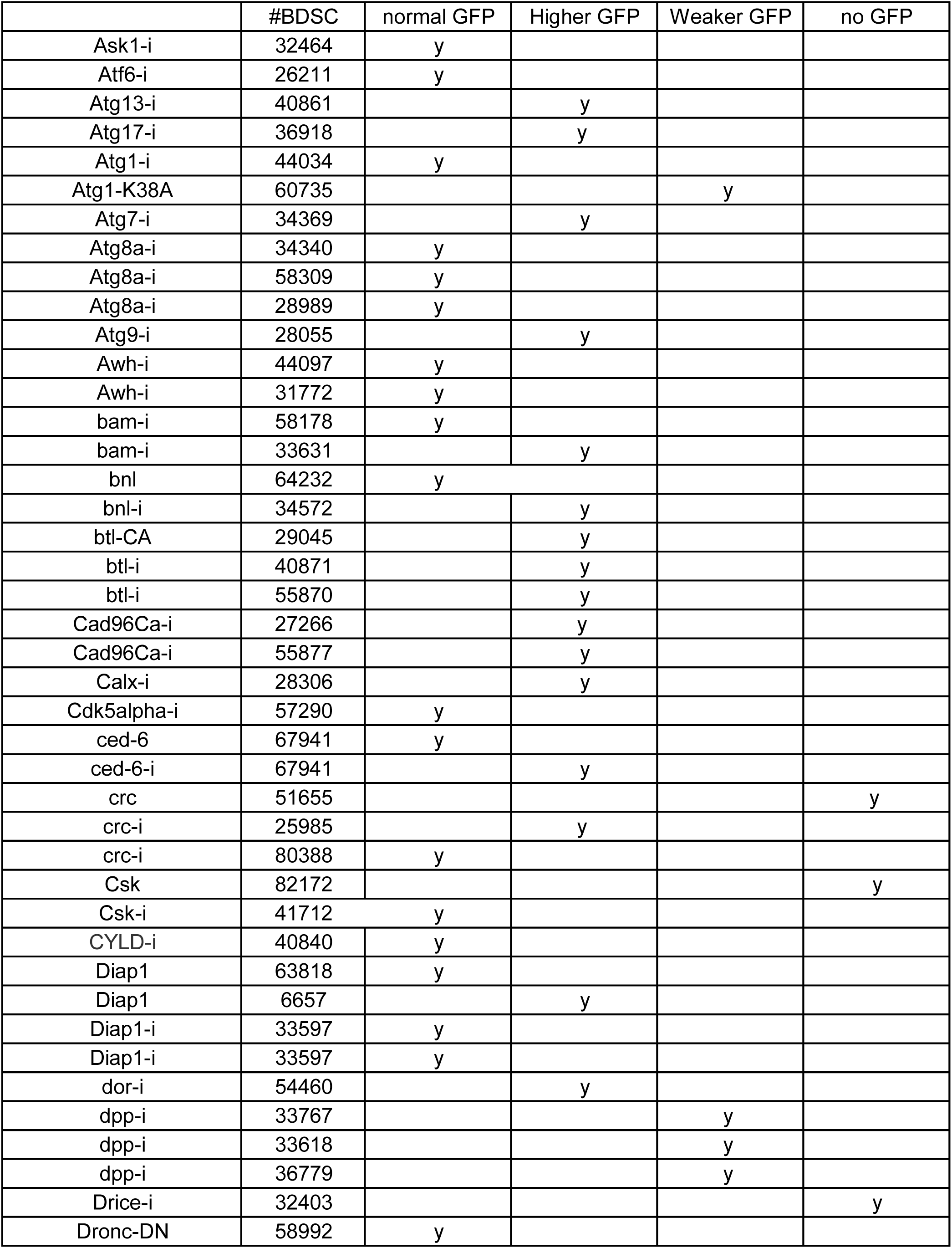
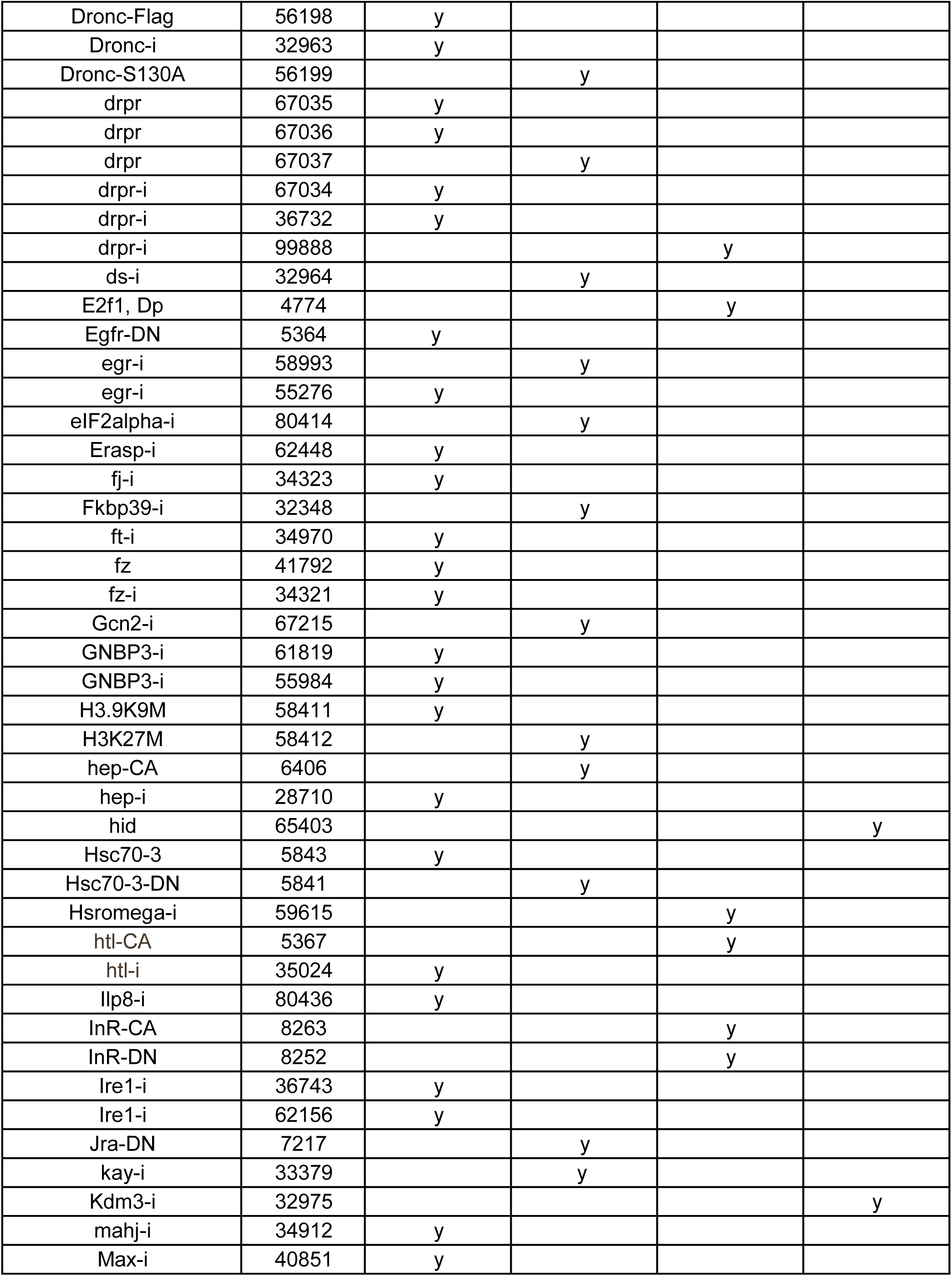
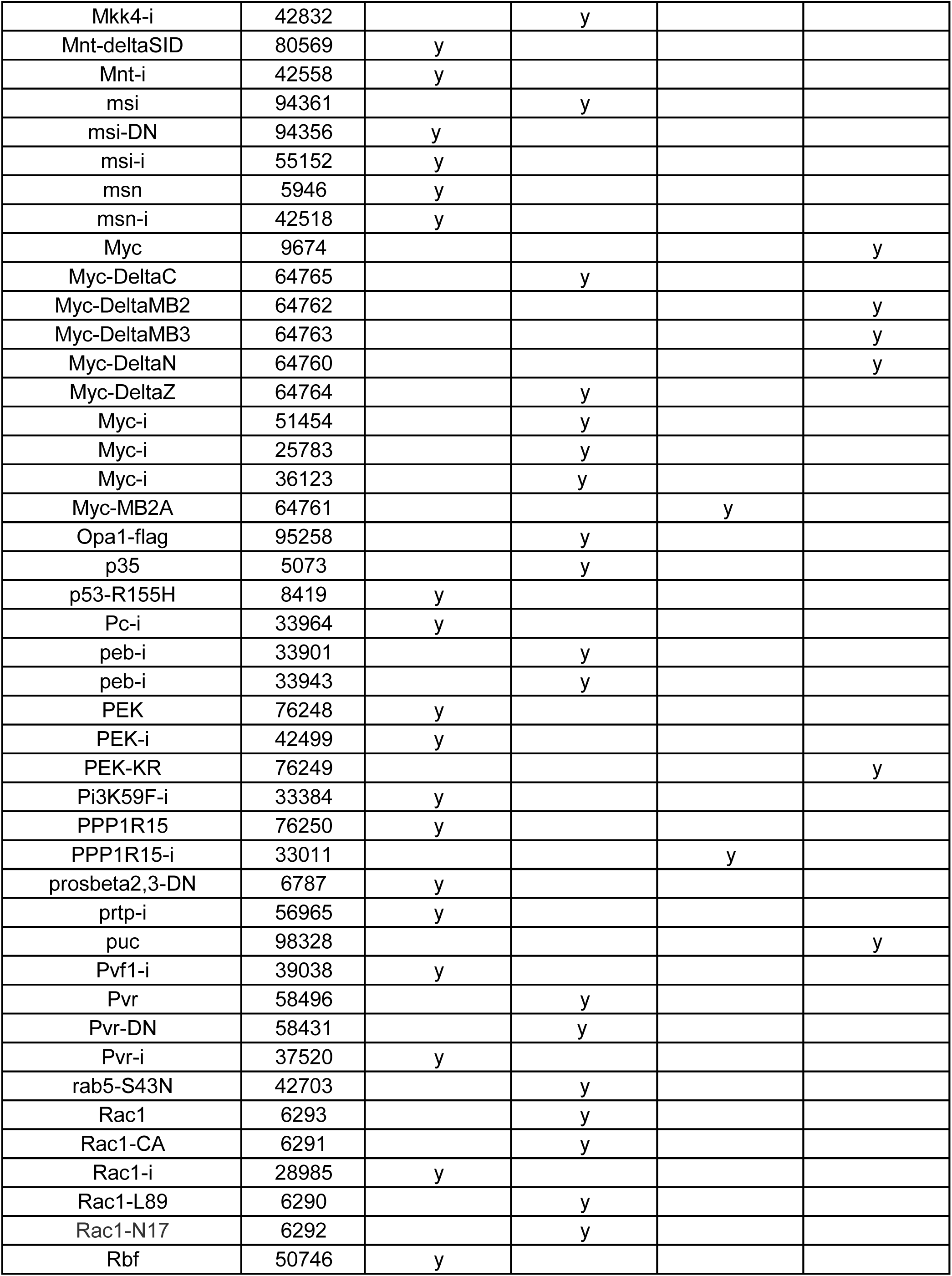
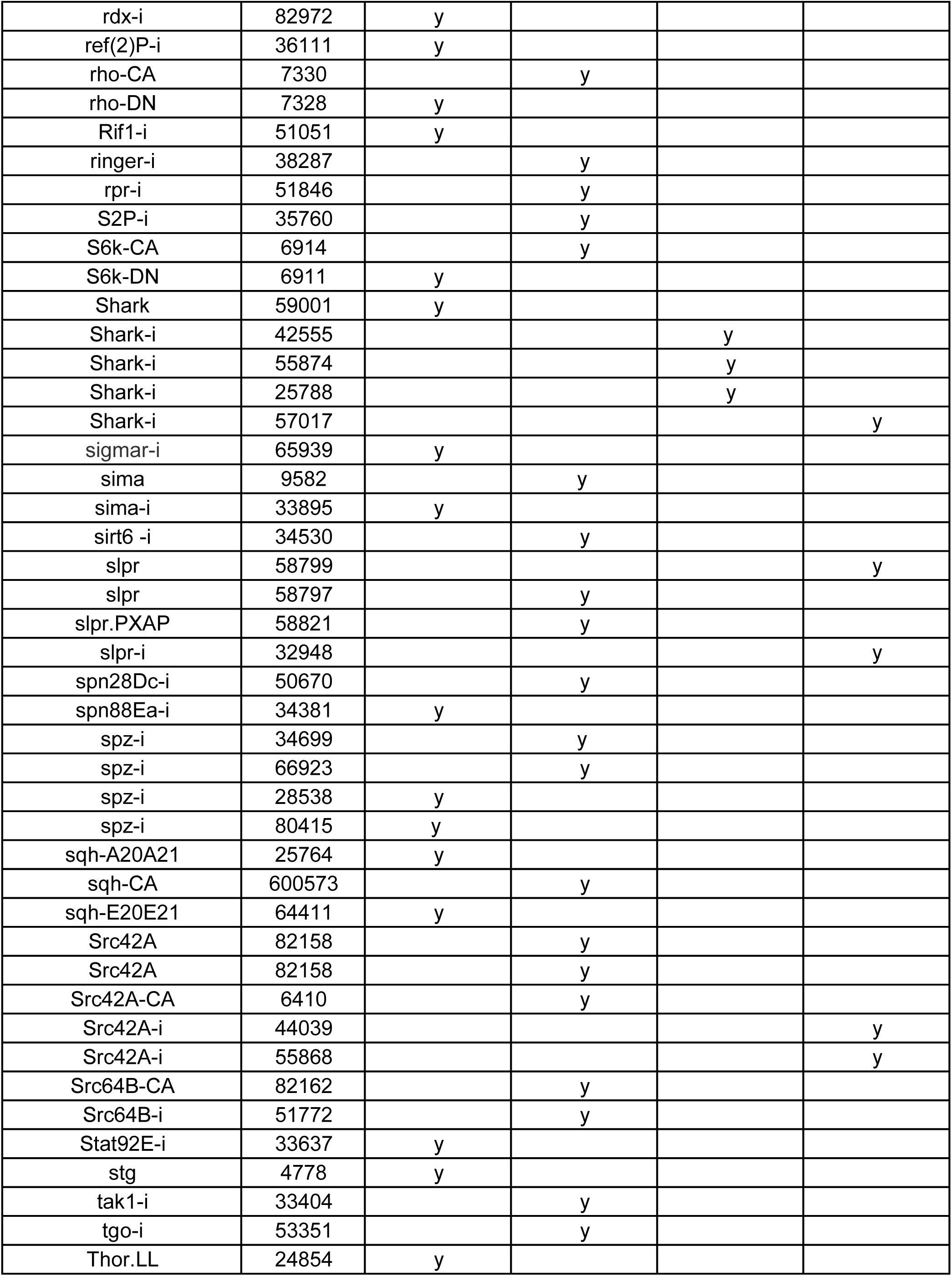
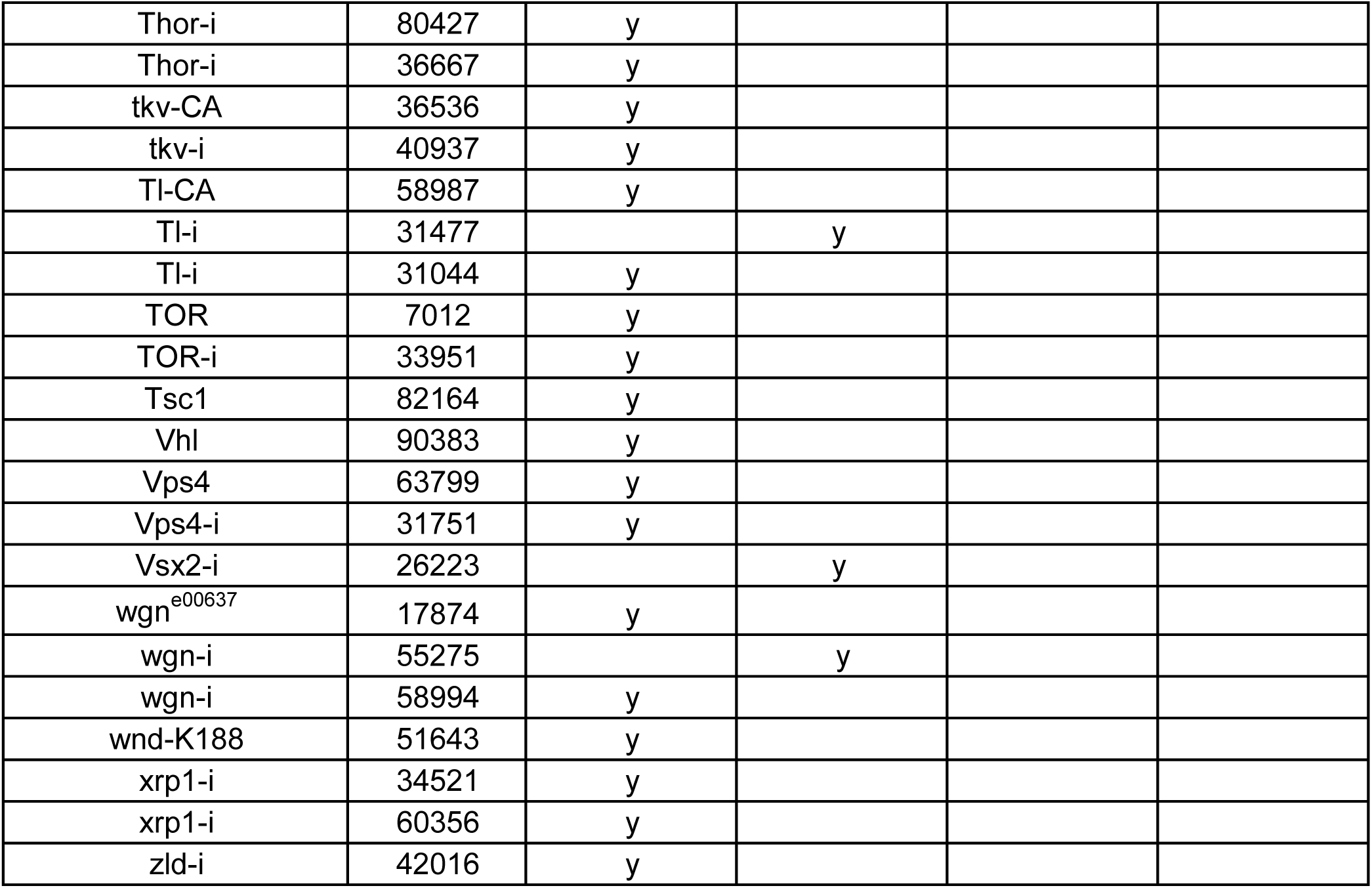
List of fly strains used in the genetic screen.

**Table S2:**
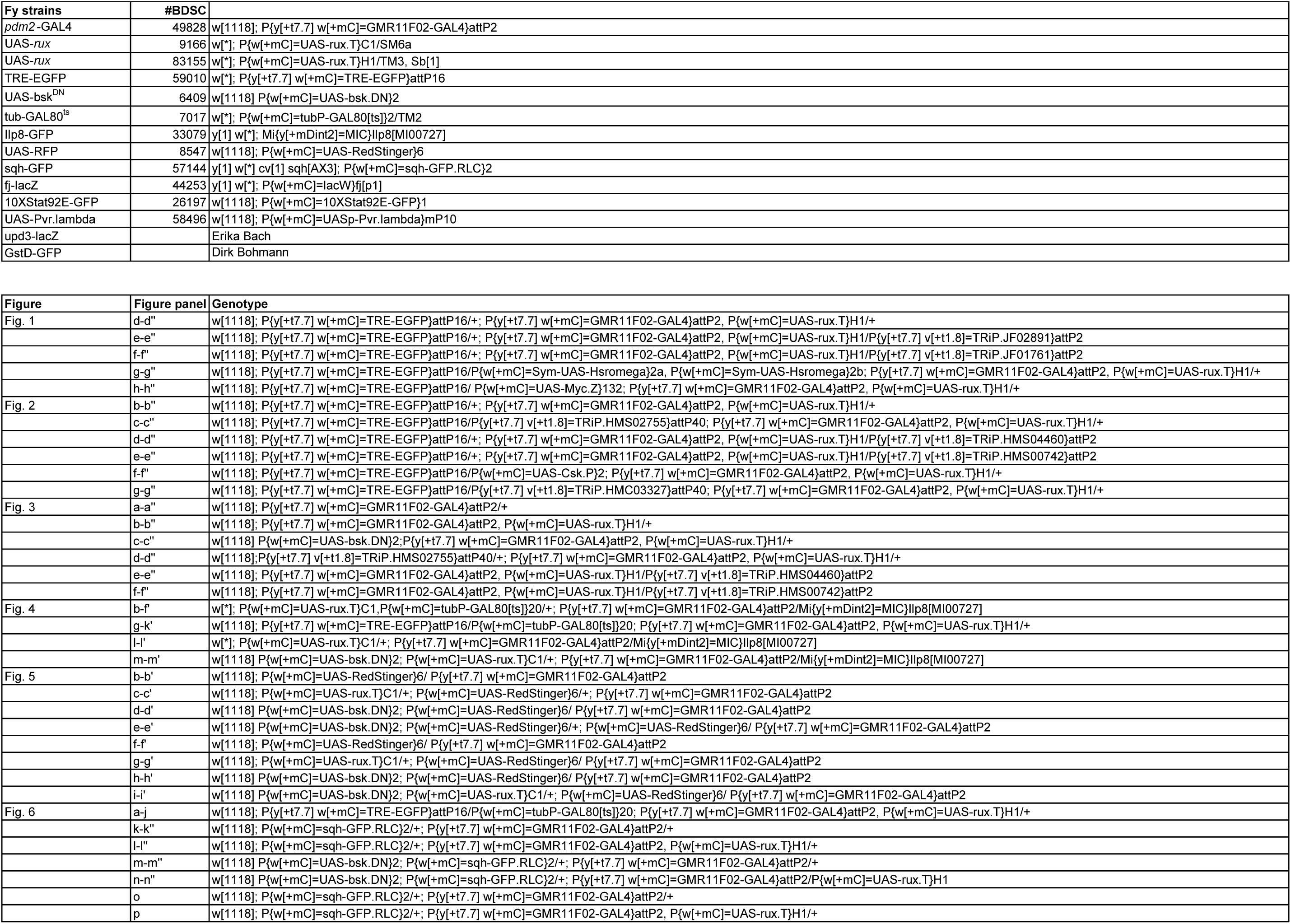
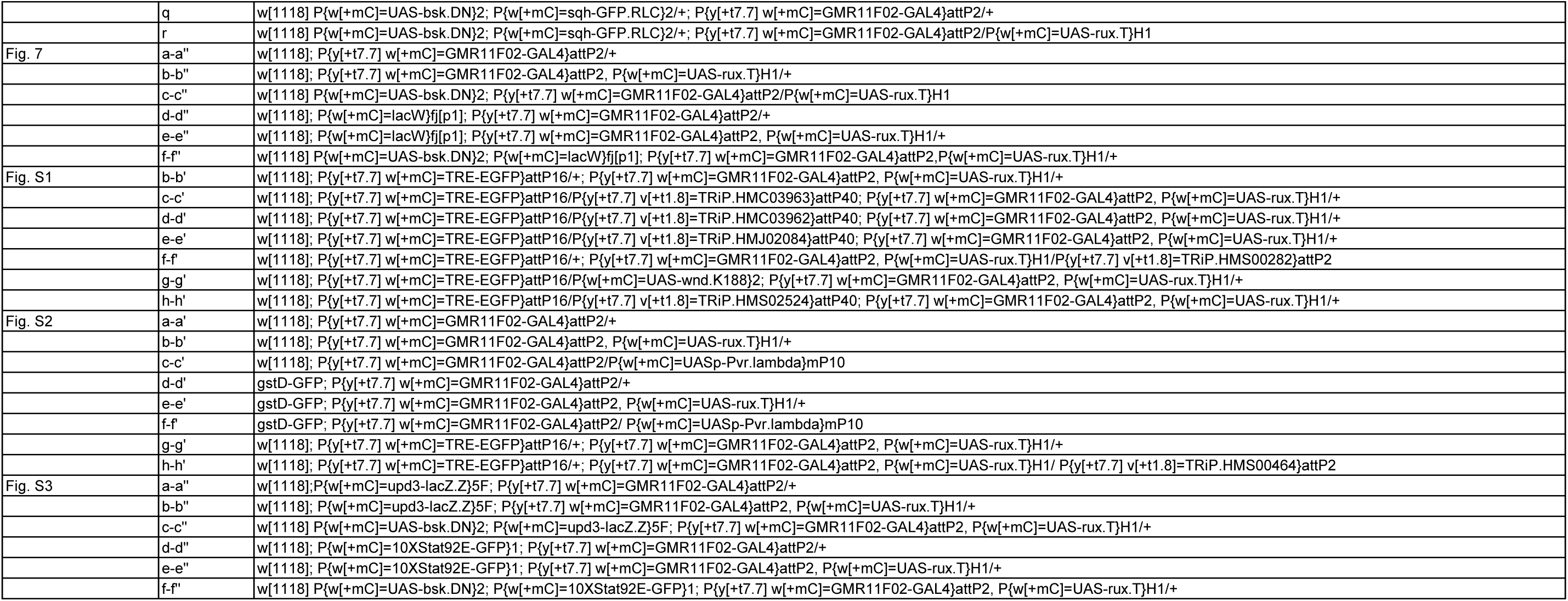
Other reagents and fly strains used.

Video S1: The morphology of a wing disc with *pdm2*-GAL4>*mRFP* expression.

Video S2: The morphology of a wing disc with *pdm2-GAL4*>*mRFP*, *rux* expression.

Video S3: The morphology of a wing disc with *pdm2-GAL4*>*mRFP*, *bsk^DN^* expression.

Video S4: The morphology of a wing disc with *pdm2-GAL4*>*mRFP*, *bsk^DN^, rux* expression.

